# Conserved 3’ Stem-Loop Structures Enable Comprehensive Analysis of Bacterial Transcription Termination in Metagenomes

**DOI:** 10.1101/2023.10.02.560326

**Authors:** Yunfan Jin, Jiyun Cui, Hongli Ma, Fei Gan, Zhenjiang Zech Xu, Zhi John Lu

**Affiliations:** MOE Key Laboratory of Bioinformatics, Center for Synthetic and Systems Biology, School of Life Sciences, Tsinghua University, Beijing 100084, China; Institute for Precision Medicine, Tsinghua University, Beijing 100084, China; Hubei Key Laboratory of Cell Homeostasis, College of Life Science, TaiKang Center for Life and Medical Sciences, Wuhan University, Wuhan, 430072, China; School of Mathematics, Harbin Institute of Technology, Harbin 150001, China; State Key Laboratory of Food Science and Resources, Nanchang University, Nanchang 330047, China; School of Mathematics and Computer Sciences, Nanchang University

**Keywords:** bacteria terminator, transcription termination, transcription unit prediction

## Abstract

Bacterial transcription termination is a critical yet underexplored mechanism of gene regulation in microbial ecosystems. Existing computational tools, however, primarily focus on predicting transcript 3’ ends generated by Rho-independent terminators (RITs) in model species, leaving significant gaps in understanding those generated by Rho-dependent terminators (RDTs), especially in non-model species. To address these limitations, we developed BATTER (BActeria Transcript Three Prime End Recognizer), a comprehensive computational tool for bacterial transcript 3’ termini prediction. BATTER builds on the observation that conserved stem-loop structures are frequently associated with 3’ ends of primary transcripts generated by both RIT and RDT mechanisms across distantly related bacterial species. BATTER demonstrated its advantage compared to existing tools. It enabled a comprehensive analysis of 42,905 representative bacterial genomes, uncovering that stem-loop structures exhibit clade-specific properties with greater variations between species than between gene families. Notably, BATTER uncovered that certain *Cyanobacteria* lineages, despite lacking Rho homologs, harbor Rho utilization (RUT) site-like sequences near 3’ ends, with preliminary experimental validation in *E. coli* suggesting their partial functionality in transcription termination. Additionally, BATTER systematically identified pervasive premature termination events in antimicrobial resistance (AMR) genes, highlighting their regulatory roles in translation protection and drug efflux. This study advances our understanding of transcription termination across diverse bacterial lineages and provides a robust computational approach for exploring transcription regulation in complex microbial ecosystems.

## Introduction

Bacterial transcription terminators are RNA elements that dissociate RNA polymerase complexes and induce transcription termination. They are traditionally classified into two categories: Rho-independent terminators (RITs) and Rho-dependent terminators (RDTs)^[1, 2]^. In model species *Escherichia coli* and *Bacillus subtilis*, RITs are GC-rich RNA hairpins followed by a stretch of uracils. RDTs typically contain a ~60 nt subsequence called Rho utilization (RUT) sites, which are the primary binding sites of the Rho protein^[3]^. RUT sites are rich in YC (a pyrimidine followed by a cytosine) dimers that are regularly spaced by distances of around 10 nt^[2, 4]^. They are also characterized by features such as high C/G ratios (referred to as C>G “bubbles” ^[3]^) and depletion of secondary structures^[2]^.

RDTs were reported to account for 20%-30% of the primary transcription termination events in *E. coli* ^[5]^, but play a less important role in *B. subtilis*^[6]^. Computational prediction algorithms for RIT have been established in *E. coli* and *B. subtilis*^[7-9]^, yet predicting RDTs remains challenging. This difficulty arises because the known sequence patterns of RDTs are less compact and distinct than those of RITs, even in well-studied model organisms ^[2, 3, 10]^. Available tools predict a paucity of RITs in some bacterial species^[7, 9]^, raising questions as to whether these species mainly use Rho-dependent mechanisms or their RITs have alternative sequence patterns. This gap in computational predictions poses a significant challenge for studying transcription termination mechanisms across bacterial lineages, especially in metagenomic datasets where diverse bacterial species are often present. Accurate terminator prediction is essential for understanding gene regulation and community dynamics in these complex microbial environments. Limitations in current computational methods thus present a major barrier to comparative analyses of transcription terminators across diverse bacterial clades, impeding insights into transcriptional regulation in diverse microbial communities.

Recently, several high-throughput 3’ ends mapping experimental techniques, such as term-seq^[11]^, Rend-Seq^[12]^, and SEnd-seq^[13]^ were developed. These techniques detect 3’ ends of RNAs, regardless of whether the transcripts employ RITs or RDTs for transcription termination. Although a 3’ ends mapping experiment alone cannot determine whether a transcript end is generated by RIT or RDT mechanism, the rough classification can be achieved by evaluating changes in RNA-seq coverage flanking detected transcript 3’ ends upon inactivation of Rho protein by gene knockout^[14]^, inducible protein depletion^[15]^ or the small molecule inhibitor BCM (bicyclomycin)^[16, 17]^. Available term-seq data suggested stem loop structures are pervasively associated with transcript 3’ ends in bacteria species belonging to *Proteobacteria, Firmicutes, Actinobacteria, Spirochaetota*, and *Cyanobacteria* lineages^[12, 17-26]^. While RUT sites are thought to be depleted of RNA secondary structure^[1, 2]^, based on 3’ end mapping techniques, it was reported that in *E. coli*, transcript ends generated by the Rho-dependent mechanism can fold into stem loop structures reminiscent of that in RITs with comparable folding energies, but typically lack the U-rich tracts^[5]^. In 3’ end mapping data of *Streptomyces lividans* TK24, stem loop structures are also frequently observed in 3’ ends that are not associated with U-rich sequences^[19]^. A possible explanation for this apparent discrepancy is that, for transcripts terminated by RDTs, 3’ ends are rapidly degraded by intracellular exoribonuclease such as PNPase after transcription termination, until reaching the stem loop structures which prevent further degradation^[5, 27]^.

The stem loop structures are crucial for maintaining transcript integrity; as a result, for functional transcripts, a 3’ end-associated stem loop should be positively selected in evolution^[5]^. If this property is broadly conserved in bacteria, as a stem loop structure is a stronger feature compared to sequence patterns of RUT sites, this observation can be leveraged for comprehensive prediction of 3’ ends of primary transcript in diverse bacterial lineages, regardless of Rho factor dependency. The steady-state 3’ ends do not correspond to real transcription termination sites^[28]^; and it is worth noting that in addition to terminating transcription at the end of transcription units, an important function of Rho protein is suppressing pervasive transcription, and the distribution of RUT sites are not restricted to the ends of transcription unit, but pervasive across the genome, especially in antisense of CDSs^[3, 29]^. These pervasive transcription products are not as stable as primary transcripts and often elude detection of term-seq^[30]^. However, for applications like operon prediction and noncoding RNA (ncRNA) prediction, predicting stable 3’ ends of primary transcripts is sufficient.

In this study, through term-seq data analysis, we validated that at least for primary transcription terminations, 3’ ends generated by both RITs and RDTs are frequently associated with conserved stem loop structures in diverse bacterial species. Based on this observation, exploiting advancements in both experimental data and computational techniques, we present BATTER, a computational tool for bacteria transcription termination analysis. BATTER is composed of two modules: a machine learning model based on BERT-CRF architecture for predicting steady-state RNA 3’ ends (BATTER-TPE, Three Prime End) and an accessory rule-based algorithm for detecting putative RUT sites (BATTER-RUT). Systematic evaluation on diverse metrics suggested BATTER outperformed existing tools for both 3’ ends detection and RUT sites prediction.

Using BATTER, we comprehensively predicted transcript 3’ ends and putative RUT sites near the predicted 3’ ends in representative bacterial genomes. We characterized the variations of these conserved stem loops across diverse bacterial lineages and evaluated their coevolution with the presence of Rho proteins and putative RUT sites. Our analysis suggests the prevalence of RDT and RIT is negatively correlated in different species. Unexpectedly, we also detected RUT sites-like signatures in *Cyanobacteria* species, which lack Rho homologs. This signature is correlated with more recently evolved traits. Preliminary experimental analysis suggested some of these RUT site-like sequences are capable of inducing transcription terminations in *E. coli*. As another application, by predicting premature terminations of known antimicrobial resistance (AMR) genes, we systematically identified premature termination-based regulations and related ncRNAs. Our analysis suggests that premature termination-based regulations are pervasive in AMR genes related to translation protection and drug efflux.

## Methods

### Curation of experimentally supported 3’ ends

Known transcript 3’ ends with experimental supports used in this work were curated from previous studies (Table S1). The transcript 3’ ends of the metaterm-seq dataset^[11]^, which are not provided by the original publication, were identified by a customized pipeline. In brief, metaterm-seq reads and RNA-seq reads of 2 human oral metagenome samples were downloaded from PRJEB12568. Metaphlan2^[31]^ was used for species profiling; species with abundance > 1% in at least one sample (either RNA-seq or term-seq) were considered for downstream analysis. Reads were then mapped to corresponding reference genomes downloaded from genbank (Table S3). Coverage of the captured 3’ ends was summarized with HTSeq^[32]^. Local max positions within 50 nt windows covered by at least 6 reads are considered as transcripts’ 3’ ends with experimental support.

### Identification of Rho-dependent regions

The RNA-seq reads of *E. coli* (GSE41939), *B. subtilis* (GSE195579), and *B. burgdorferi* (GSE222085) were mapped to the corresponding reference genomes (Table S2) with bowtie2^[33]^ using the default parameters. For *M. tuberculosis* data (E-MTAB-11753), bam files were downloaded from http://ftp.ebi.ac.uk/biostudies/fire/E-MTAB-/753/E-MTAB-11753/Files. The identification of Rho-dependent regions was implemented using a customized python script. In brief, using a stride of 50 nt, for each considered genome location, the numbers of RNA-seq fragments assigned within the upstream 200 nt (*U*_200_) and the downstream 200 nt (*D*_200_) were summarized, replicates were combined by summation, and the ratio *D*_200_/*U*_200_ measures the level of read-through at this location. Single-sided Fisher’s exact test was used to evaluate whether the read-through level in Rho protein inhibition group was greater than the control group. For *B. subtilis*, data from wild type (WT) and Rho knockout strains in the exponential growth stage were considered. For *M. tuberculosis*, samples at 6h after Rho protein depletion were used. Regions with odds ratio > 3 and *P*-value < 0.001 were classified as Rho-dependent regions, those with odds ratio < 1 were classified as Rho-independent regions, and the remaining ones were classified as “others”. 3’ ends that overlap with Rho-dependent regions, Rho-independent regions, and “others” are categorized as RDT, RIT, and unclassified, respectively. The folding energy of upstream 45 nt, and downstream 5 nt sequences flanking annotated 3’ ends were evaluated with RNAfold utility in ViennaRNA package^[34]^.

### Implementation and benchmarking of the BATTER algorithm

See Supplementary Information for details of the implementation.

### Calculation of Rho dependency scores

To quantify the contribution of Rho-dependent termination, for BATTER-TPE predicted primary transcript 3’ ends at an FPR cutoff of 0.1/KB, the upstream 100nt and downstream 300nt sequences are extracted and scanned by BATTER-TPE with default parameters. For each 3’ end, the best-scoring RUT site is extracted. If no RUT site is predicted, a score of 0 is assigned. The average score across all primary 3’ ends was utilized as a measure of Rho dependency.

### *Cyanobacteria* traits analysis

For *Cyanobacteria* traits analysis, the trait data were curated based on several previous studies^[35-37]^. *Cyanobacteria* genomes with PsaA2, PsaB2, and PsbA4 gene orthologs were annotated as FarLip, and genomes with apcD4, apcB3, and isiX gene orthologs were annotated as LoLip. Species with complete genomes were considered. The species tree is built with FastTree^[38]^ using multiple sequence alignment of 120 universal marker genes defined by gtdb-tk^[39]^.

### Fluorescence reporter assay

All cloning was performed in *E. coli* DH5α strain. Transformed cells were cultured in LB medium and on 1.5% (w/v) LB agar plates supplemented with 15 μg/mL chloramphenicol as required. Sequences encoding sfGFP, the rrnB T1 terminator, and mCherry were assembled into a vector backbone containing the replication origin of pSC101 to yield recombinant plasmid pXG10SFM. To generate the negative control plasmid pXG10SFM0, the rrnB T1 terminator of pXG10SFM was replaced with a 96 bp-region from the lacI coding sequence. To construct plasmids containing predicted RUT sites from the nitrogen-fixation gene cluster in the cyanobacterium *Nostoc sp*. PCC 7120 and FaRLiP and LoLiP gene clusters in *Chlorogloeopsis fritschii* PCC 6912, pXG10SFM was first digested with XhoI and KpnI (Thermo Fisher Scientific). RUT site-like sequences were generated by PCR using two synthesized oligonucleotide primers, and the amplicons were assembled into the digested pXG10SFM, followed by transformation into *E. coli* DH5α. Transformants were verified by DNA sequencing. The recombinant plasmids and primers used in this study are listed in Table S6 and Table S7.

To measure fluorescence in *E. coli*, transformants were first inoculated into 1 mL LB medium supplemented with 15 μg/mL chloramphenicol in 24 well-plates and grown overnight at 37°C with constant shaking at 200 rpm. Cells were washed and resuspended in PBS buffer (10 mM Phosphate, 137 mM NaCl, 2.7 mM KCl, pH=7.4). To initiate the assay, samples were diluted 1:10 into a 96-well Black/Clear bottom microplate to a final volume of 200 μL. Culture density (OD600) and fluorescence were measured by a multi-mode microplate reader Cytation3 (BioTek). Fluorescence intensities of sfGFP and mCherry were read at excitation/emission wavelengths 488 nm/515 nm and 579 nm/616 nm, respectively.

### Scanning premature terminators of AMR genes

AMR genes in GEMs’ representative genomes were annotated with AMRfinder^[40]^. 800 nt sequences (upstream 400 nt and downstream 400 nt) flanking start codons of detected AMR genes are subjected to terminator scanning. Predicted terminators located within the upstream 128 nt and downstream 16 nt of start codons are retained. These upstream 128 nt and downstream 16 nt terminator-containing sequences were grouped by AMR families and searched for conserved motifs with CMfinder^[41]^. The secondary structural covariations of resulting motifs were evaluated with R-scape^[42]^.

## Results

### A conserved stem loop is associated with the transcript 3’ end regardless of the factor dependency

To inspect the secondary structures of detected 3’ ends, we curated available 3’ ends annotations determined by term-seq, and RNA-seq data with and without Rho inhibition from 4 phylogenetically diverse species (belonging to 4 distinct phyla), including *Escherichia coli* ^[5, 16]^, *Bacillus subtilis*^[14, 27]^, *Borrelia burgdorferi*^[17]^ and *Mycobacterium tuberculosis*^[15]^(Table S1, Table S2). It has been reported RNA-seq is more effective in identifying Rho-dependent regions compared to term-seq, as term-seq tends to detect steady-state 3’ ends instead of RUT sites^[30]^. Using RNA-seq data, the detected 3’ ends were classified as RIT or RDT according to changes in read-through upon Rho inhibition (see Methods). The distances between the 3’ ends defined by term-seq and their closest upstream stop codon peaked at around 50 nt (Figure S1 A). 76.5% and 82.9% 3’ ends have distances less than 100 nt and 150 nt, respectively, and a cutoff of 128 nt should cover the majority of primary transcription termination events. For this reason, here we define “primary” termination sites as transcript 3’ ends located within 128 nt downstream of the CDSs. In all four species, 3’ ends detected by term-seq exhibit lower folding energies compared to the genomic background, particularly at primary termination sites. Notably, the folding energies of primary 3’ ends assigned to RDTs and those assigned to RITs are similar (Figure 1A).

**Figure 1.**
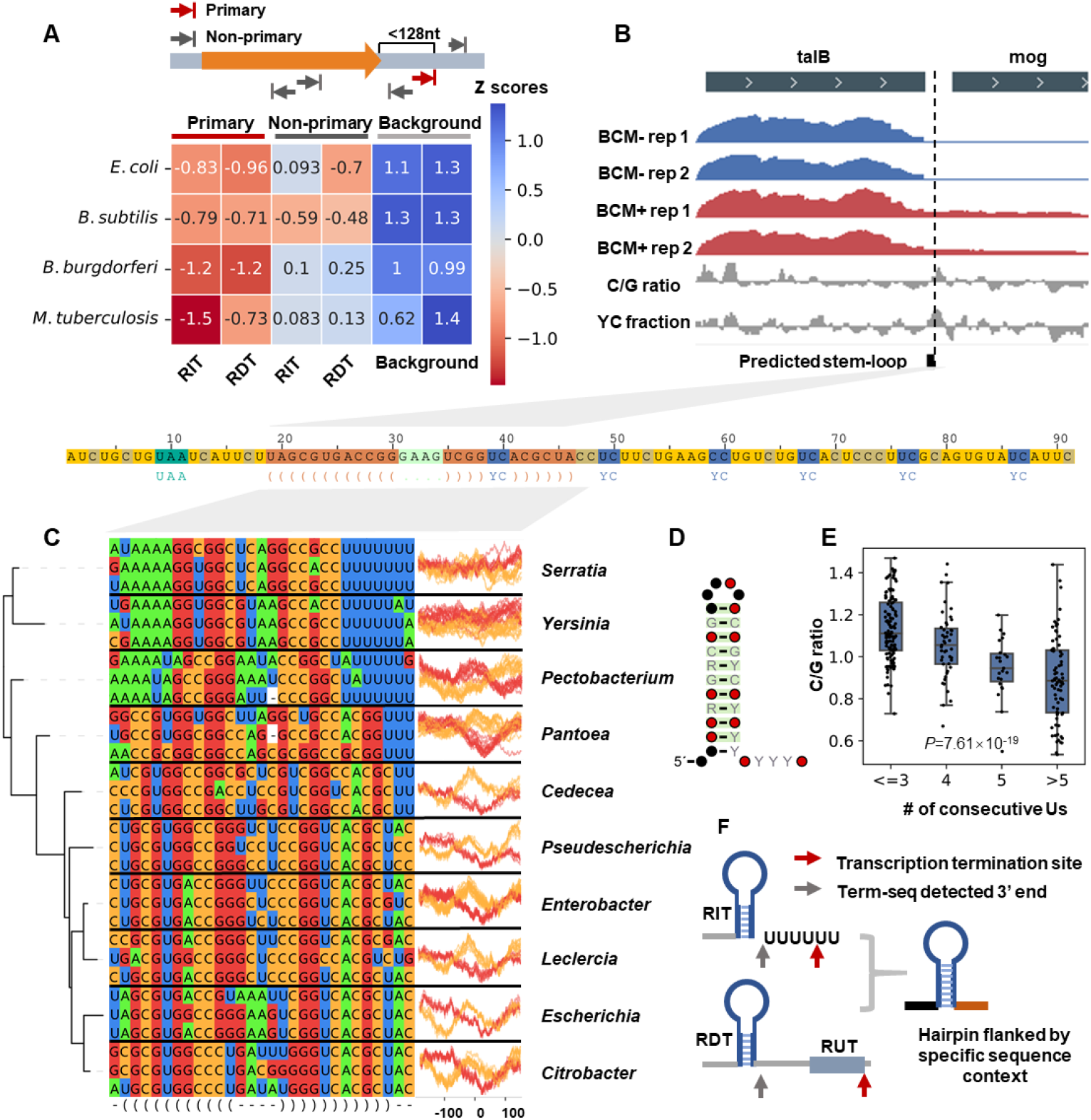
A conserved stem loop is associated with transcript 3’ end regardless of the Rho factor dependency A. Z scores of average folding energy in 4 distantly related species, stratified by Rho factor dependency (RITs and RDTs) and genomic context (primary 3’ ends, non-primary 3’ ends, and background) B. RNA-seq coverage near the 3’ end of E. coli talB under BCM treatment (BCM+ rep 1, BCM+ rep 2) and control (BCM-rep 1, BCM-rep 2) conditions. Local C/G ratio and density of YC dimers are also shown. C. Multiple sequence alignment (MSA) of the conserved stem loop structure in downstream sequences of talB gene in the *Enterobacteriaceae* family. For each genus, sequences from 3 representative species are shown. In the right panel, for all species of each genus in GEMs representative genomes, G ratios and C ratios (labeled in red and orange, respectively) of 101 nt sequences centered at each nucleotide from 128 nt upstream to 128 nt downstream relative to ends of the stem loops are shown. D. Secondary structure covariation of the identified stem loop. Base pairs with R-scape E value < 0.05 are labeled in green, and nucleotides with sequence identity > 97% and > 90% are labeled in red and black, respectively. R indicates purine, Y indicates pyrimidine. E. Correlation between maximum lengths of consecutive uracils (Us) and C/G ratio within 128 nt sequences flanking ends of the stem loops downstream talB in all considered Enterobacteriaceae genomes. F. Schematic illustration of common and distinctive features of RITs and RDTs. For RITs, term-seq sites indicate the location of terminator stem loops, while for RDTs, term-seq sites indicate stem loops upstream of real termination sites. Both RITs and RDTs are characterized by stem loops flanked by particular sequence contexts.

As an example, using data from a previous study ^[16]^, the RNA-seq read coverages near 3’ end of *E. coli* talB gene were illustrated (Figure 1B). Upon BCM treatment, talB has a higher level of transcription read-through. The presence of regularly spaced YC dimers and a high C/G ratio confirmed that the Rho-dependent mechanism is involved. There is a stem loop structure immediately upstream of its 3’ end. To analyze the conservation of the stem loop, we identified homologs of talB gene within taxa *Enterobacteriaceae* in representative genomes of GEMs (Genomic catalog of Earth’s Microbiomes) dataset ^[43]^ and searched conserved hairpins near stop codons (upstream 16 nt and downstream 128 nt) with CMfinder^[41]^. As shown in Figure 1C, species within *Escherichia* genus, as well as *Citrobacter, Leclercia, Enterobacter*, and *Pseudescherichia*, exhibit RDT-like properties, including the presence of C>G bubbles and the absence of U-rich tracts, while in *Serratia, Yersinia*, and *Pectobacterium* species, the homolog transcripts are ended with a typical RIT-like motif, characterized by a stretch of uracils (Us) and lacking a C>G bubble in downstream sequences. R-scape^[42]^ analysis revealed strong covariation of the predicted base pairing (Figure 1D). We also observed a significant negative correlation between the maximum length of U runs near the stem loops and the C/G ratio within 100 nt upstream and 100 nt downstream flanking sequences (Figure 1E, ANOVA *P*=7.61×10^-19^). These results suggested the hairpin is structurally conserved in all homolog transcripts, despite the apparent differences in Rho factor dependency. talB is not a special case, and similar phenomena are observed in other genes; 3 of such genes were exemplified in Figure S1 B-C.

Taken together, primary transcription terminations conducted by both RIT and RDT are characterized by conserved stem loops, and the stem loop structures can be more conserved than Rho factor dependency. This feature is likely to be conserved in diverse bacterial lineages. Thus, it is possible to predict transcript 3’ ends generated by both RITs and RDTs in a common framework, that is, predicting stem loop structures embedded in certain sequence contexts (Figure 1F).

### Models for transcript 3’ ends and RUT sites prediction

We collected transcript 3’ ends defined by published genome-wide studies^[12, 18-26]^ (Table S1). Naively training models on experimentally verified 3’ ends presented several challenges, including phylogeny distribution bias (Figure S2 B), limited sample size, and variations due to different experimental settings. Based on the conservation of primary 3’ ends associated stem loop structures, we developed a data augmentation pipeline (Figure S2 A) and generated ~2.5 million putative 3’ ends associated stem loop instances (Figure S2 C). Then we trained a BERT-CRF model called BATTER-TPE, for bacterial transcript 3’ ends prediction (Figure S2 D-E). While BATTER-TPE predicts the 3’ ends of primary transcripts, it cannot locate potential RUT sites around these 3’ ends. To address this, we also developed a rule-based post-processing module, BATTER-RUT, which identifies putative Rho binding sites by searching YC dimers with optimal spacing (Figure S3). See Methods for the implementation details of the data augmentation pipeline, BATTER-TPE, and BATTER-RUT.

We further benchmarked the performance of BATTER-TPE and BATTER-RUT, and compared them with 4 existing tools on diverse metrics. For primary transcription termination, at a false positive rate (FPR) cutoff of 0.1 per kilobase, stem loop prediction-based methods outperform Rho binding sites prediction-based methods by a large margin, even for RDT-associated 3’ ends (Figure 2A). This is within expectation, as features like a high local C/G ratio and the presence of regularly spaced YC dimers are pervasive in random sequences. For both RDT- and RIT-associated primary 3’ ends, BATTER-TPE outperforms transterm-HP and RNIE. As shown in Figure S4 A, stem-loop prediction methods, including BATTER-TPE, capture a subset of non-primary 3’ ends in *E. coli* (recall of around 40%) and *B. subtilis* (recall of around 90% for RIT, 75% for RDT), but perform poorly on *M. tuberculosis* and *B. burgdorferi* (recall below 10%). Nonetheless, these stem-loop-based approaches generally yield better or comparable results to RUT site-based prediction methods. This aligns with our observation that folding energy drops near non-primary transcript ends are less pronounced than at primary transcript ends, indicating that non-primary termination events may involve complex, clade-specific properties.

**Figure 2.**
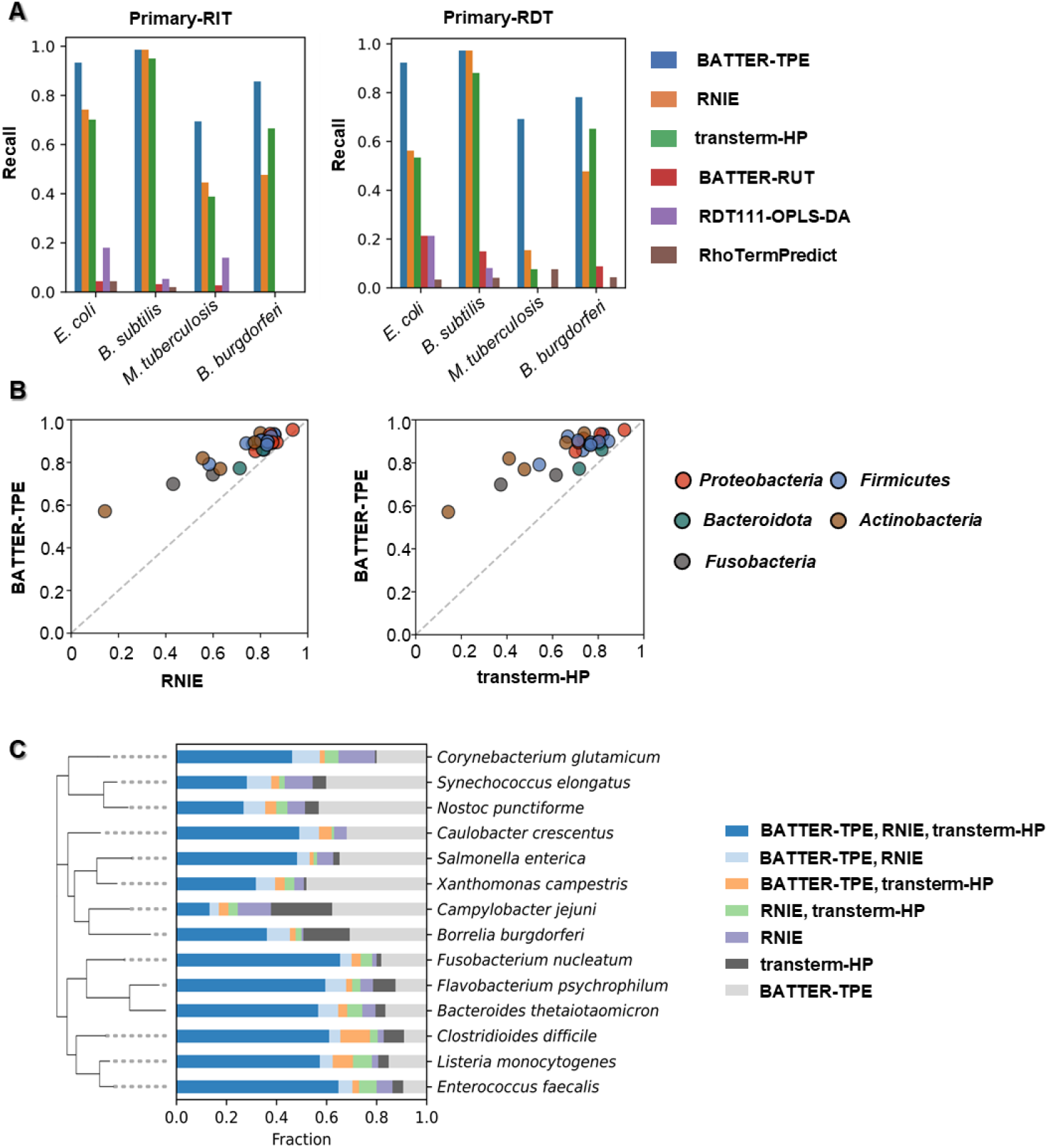
Performance evaluation of the 3’ end prediction model A. Recalls of stem loop prediction-based methods (BATTER-TPE, RNIE, and transterm-HP) and RUT sites prediction-based methods (BATTER-RUT, RDT111-OPLS-DA, and RhoTermPredict) at FPR of 0.1/KB. B. Performance comparisons for BATTER vs. RNIE (left panel) and BATTER vs. transterm-HP (right panel) on term-seq dataset of the oral microbiome. Recalls of two methods at an FPR of 0.1/KB on different species are plotted against each other. Colors indicate different phyla Overlaps between RNIE, BATTER, and transterm-HP’s predictions on curated differential RNA-seq datasets.

The performances of BATTER-TPE and published tools on 3’ ends prediction were also evaluated on term-seq data of the human oral microbiome^[11]^. For species with abundance > 1% (Table S3), BATTER-TPE consistently detects more 3’ ends (Figure 2B). This improvement is particularly noticeable for phyla like *Actinobacteria* and *Fusobacteria*.

Since RUT site prediction-based methods performed poorly in detecting transcript 3’ ends, we focused on evaluating their effectiveness in identifying RUT sites. Given the challenge of defining precise RUT locations and the lack of comprehensive genome-wide annotation^[3]^, we used Rho-dependent regions identified by RNA-seq as positive instances, as these regions are likely enriched for RUT sites. In four species, BATTER-RUT mildly outperformed the other two methods (Figure S4 B) at an FPR cutoff of 1/KB. For all 4 species, predictions of BATTER-RUT are enriched downstream of transcript 3’ ends, peaking at around 50 nt (Figure S4 C).

For genomes that lack experimental data, here we used a previously proposed approach^[9]^: There should be transcript 3’ ends in intergenic regions between convergently transcribed (“tail to tail”) gene pairs. In a random subset of ~2000 representative bacteria genomes in GEMs, at an FPR cutoff of 0.1/KB, for all the IGRs flanked by tail-to-tail gene pairs, the fraction of IGRs with predicted 3’ ends measures model performance (Figure S5 A). BATTER-TPE consistently achieves the best performance across diverse bacterial phylogeny compared to the other two methods.

The performance of BATTER-TPE on detecting 3’ ends of ncRNAs was also evaluated. Assuming the majority of ncRNAs with TSS detected by dRNA-seq have terminators, a higher number of predicted terminators located between intergenic TSS and the next CDS would indicate better performance. We curated TSSs from 14 bacteria species that are representative of major bacteria lineages (Table S4) defined by dRNA-seq, and defined intergenic TSSs as TSSs that do not overlap with protein-coding genes, and have distances larger than 128 nt to the closest start codons. We annotated such putative 3’ ends with transterm-HP, RNIE, and BATTER-TPE at an FPR of 0.1/KB. For *Firmicutes* and *Bacteroidota*, in the predictions by the three methods, around 60% overlap. However, for *Cyanobacteria, Proteobacteria*, and *Actinobacteria* species, up to 50% of such predictions are unique to BATTER-TPE, suggesting its ability to detect terminators overlooked by transterm-HP and RNIE (Figure S5 B). Figure S5 C showcases two TSS-associated terminators uniquely predicted by BATTER-TPE that matched Rfam families, which were withheld from data augmentation.

### Inter-gene and inter-clade variations of 3’ end-associated stem loops

We next explored the variations in the stem loop structures associated with transcription 3’ ends predicted by BATTER-TPE (only including primary 3’ ends) across all the GEMs genomes. As illustrated in Figure 3A, stem loops of *Actinobacteria* are characterized by long stems and often lack downstream U-rich sequences. Stem loops of most *Firmicutes* and *Bacteroidota* show only minor deviations from typical RITs. Stem loops of *Proteobacteria* tend to be shorter and have relatively large intra-phylum variations. When only considering primary 3’ ends in the predictions, 20% to 50% of CDSs in most bacteria species are predicted to have trailing terminators, and on average, 1 to 5 terminators are predicted in every kilobase of IGR sequences (Figure 3A).

**Figure 3.**
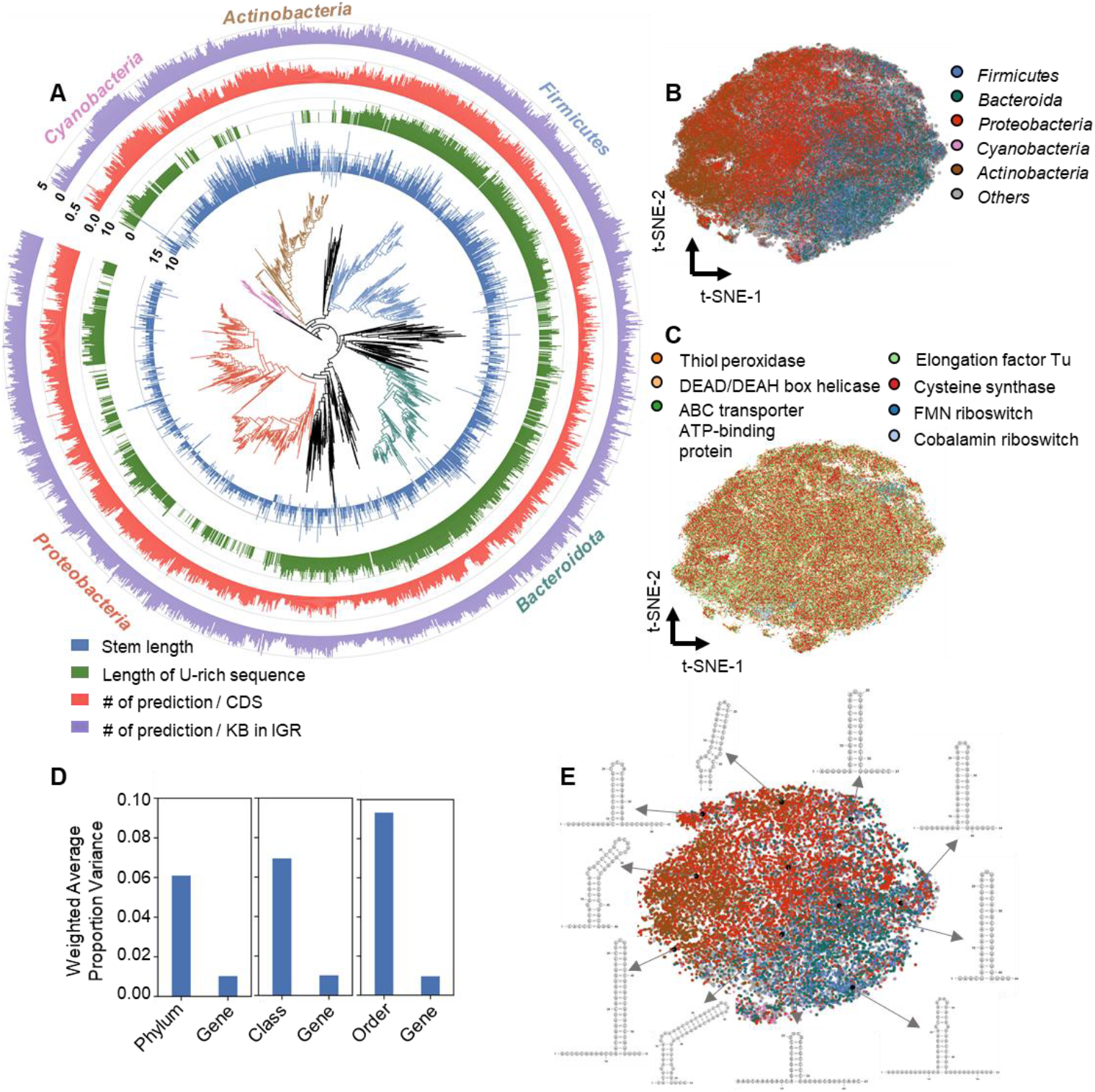
Characterization of 3’ ends associated stem loops across diverse bacteria species A. Characteristics of predicted terminators on different bacterial species. A subset of ~2000 species was sampled for visualization. From the innermost cycle to the outermost cycle, the stem lengths (blue), U-rich sequence lengths (green), number of predictions/CDS (red), and number of predictions/KB (purple) are shown. B. t-SNE (t-distributed stochastic neighbor embedding) on vector embeddings of terminators downstream of several widely distributed genes, coloring by phyla. C. t-SNE visualization same as Figure 4B, coloring by the gene families D. Weighted average variance proportion of top 36 principal components explained by different genes and different clades at the taxonomy levels of phylum, class, and order. E. Visualization of terminators associated with cysteine synthase CDS. The vector embeddings were clustered into 10 clusters with K-means, and the predicted secondary structures of cluster centers (black dots) were shown.

The 3’ end-associated stem loops show inter-gene and inter-species variations. To quantify these, we trained a neural network to project RNA sequences to numeric vectors, using a triplet loss to bring RNAs from the same Rfam family closer in embedding space while pushing RNAs from different families apart (Figure S6 A, see Methods for details). We validated the embeddings with t-SNE visualization on holdout Rfam families (Figure S6 B-C). We then analyzed terminators associated with several widely distributed genes (5 protein-coding gene families and 2 riboswitches) in all 42,905 GEMs representative bacterial genomes, projecting them into embeddings with the neural network encoder. t-SNE visualizations of the terminator embeddings (Figure 3B-C) showed that terminators of homologous genes in closely related species were similar, while in phylogenetically distant species, inter-gene variations were generally smaller than inter-species variations. To assess the contributions of taxonomy (at phylum, class, and order levels) and gene families to variations of terminators, we analyzed each of the top 36 principal components (representing ~80% of the total variance) and averaged the variance explained by each factor. As shown in Figure 3D, taxonomies consistently explained a larger fraction of variance than gene families. Taking cysteine synthase as an example, we clustered neural embeddings of associated stem loops into ten clusters using K-means and visualized the predicted secondary structures of centroid instances (Figure 3E), illustrating such cross-clade diversity of a single gene family.

### Differential contribution of Rho-dependent termination in different clades

As an application of BATTER, here we sought to quantify the relative contribution of RITs and RDTs to transcription termination in different bacterial lineages. For primary 3’ ends predicted by BATTER-TPE in each genome, we used the median length of U-rich sequences associated with these 3’ ends as a proxy to measure the contribution of RITs. We also calculated a “Rho dependency score” based on BATTER-RUT’s predictions (See Materials and Methods) to measure the contribution of RDTs. The phylogenetic distribution of Rho proteins determined by an established homolog search strategy^[35]^ was also visualized (Figure 4A). To avoid impact from genome incompleteness and potential binning error, only isolate genomes with genome completeness > 95% in GEMs representative genomes were considered.

**Figure 4.**
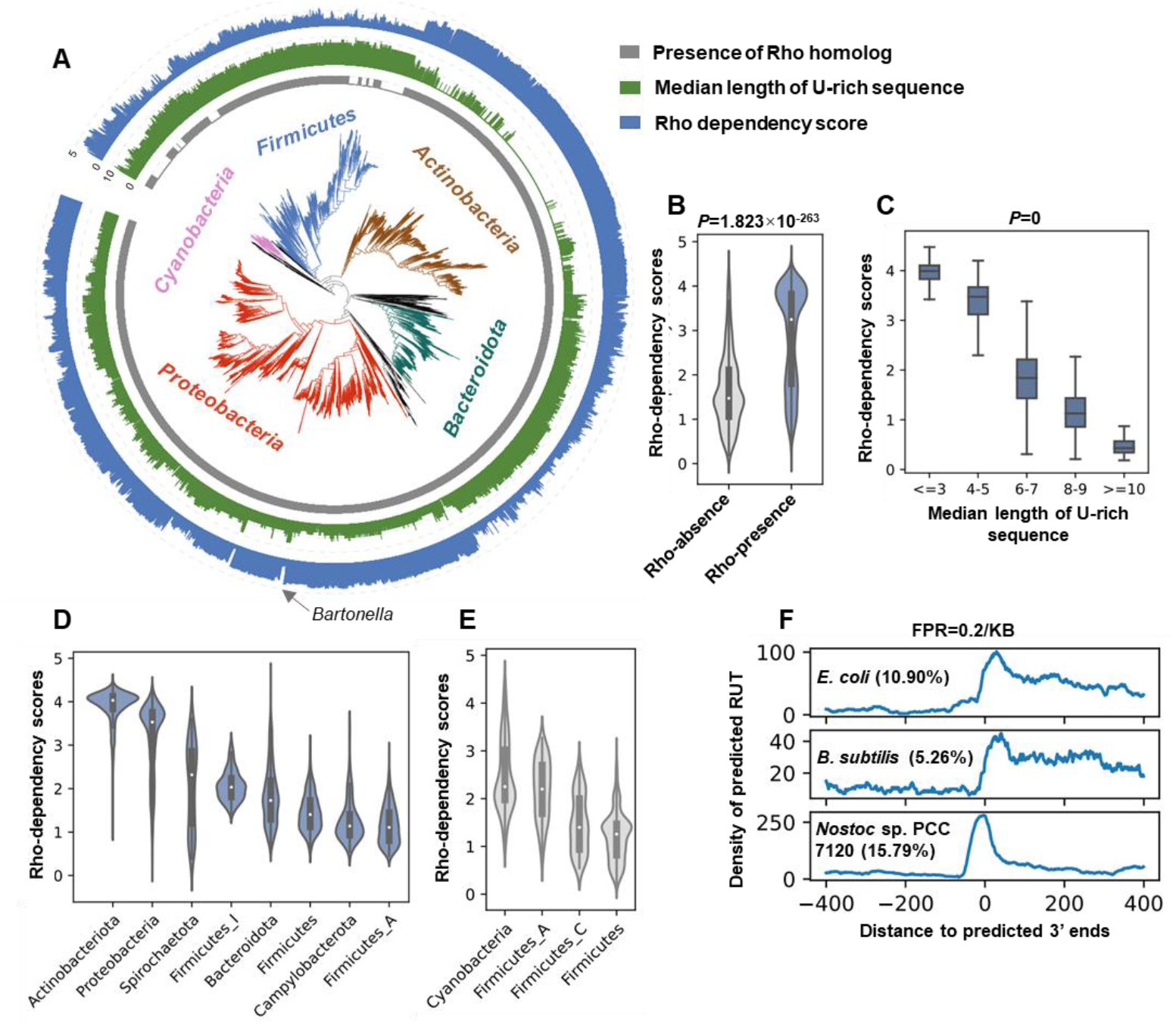
Evaluation of Rho dependency in diverse bacteria lineages A. The association between the presence of Rho protein homolog (innermost gray circle), the median length of U-rich sequences (green), and the Rho dependency score (outer circle in blue). B. Rho dependency score distribution in species with and without Rho protein homologs. *P*-value was determined by Wilcoxon test. C. The correlation between median length of U-rich sequences and Rho dependency scores. *P*-value was determined by Kruskal-Wallis test. D-E. Rho dependency score distribution in species with and without Rho protein homologs, stratified by phyla. F. Metagene plot for BATTER-RUT predicted RUT sites (FPR=0.2/KB) near 3’ ends predicted by BATTER-TPE.

In agreement with previous findings ^[35]^, we found homologs of Rho proteins are present in most of the bacterial lineages, with *Cyanobacteria* and several *Firmicutes* species as exceptions. The presence of Rho proteins and shorter U-rich sequences is strongly associated with higher Rho dependency scores (Figure 4B-C). *Actinobacteria* species have the highest average Rho dependency scores, followed by *Proteobacteria, Spirochaetota, Bacteroidota*, and *Firmicutes* (Figure 4A, Figure 4D). These findings are concordant with pervasive Rho-dependent terminations in *M. tuberculosis*^[44]^ and a diminished role of Rho protein in transcription termination in *Firmicutes* clade^[6]^. In most cases, phylogenetically related species tend to have similar Rho dependency scores. *Bartonella* species in the *Rhizobiaceae* family, which have a sharp drop in Rho dependency scores compared to other genera in the same family, is an exception. In *Bartonella* species, a basic amino acid conserved across diverse bacterial lineages (the counterpart of the R87 residue in *E. coli* Rho protein) is substituted with glutamine (Figure S7). In *E. coli*, R87 is directly involved in Rho protein-substrate interaction^[4]^, hence we hypothesized that this substitution may lead to functional deficiency, driving preference toward usage of RIT in this clade.

Taking a closer look at genomes that lack a homolog of Rho protein, surprisingly, certain *Cyanobacteria* species have unexpectedly high Rho dependency scores (Figure 4A, Figure 4E). Concurring with that, metagene plots for *Cyanobacteria* model species *Nostoc sp*. PCC 7120 suggests the predicted RUT-like sequences are also enriched downstream of predicted 3’ ends, resembling the distribution of RUT sites in species with functional Rho homologs, such as *E. coli* and *B. subtilis* (Figure 4F). However, there are notable differences: in *Cyanobacteria*, the RUT-like sequences overlap more extensively with stem loop structures, with metagene density peaks located closer to the predicted 3’ ends.

### A RUT-like signature presents in some *Cyanobacteria* species

We further investigated the phylogenetic distribution of the RUT-like sequences within *Cyanobacteria* clade and their association with traits of corresponding species. Interestingly, *Cyanobacteria* species characterized by ancient traits^[45]^—such as unicellularity and marine habitats,—tend to lack RUT-like signatures. In contrast, species with more recently evolved traits, such as the nitrogen fixation capability, far-red light photoacclimation (FarLip), and low-light photoacclimation (LoLip)^[46]^, tend to have stronger RUT-like signatures (Figure 5A-B, Table S5).

**Figure 5.**
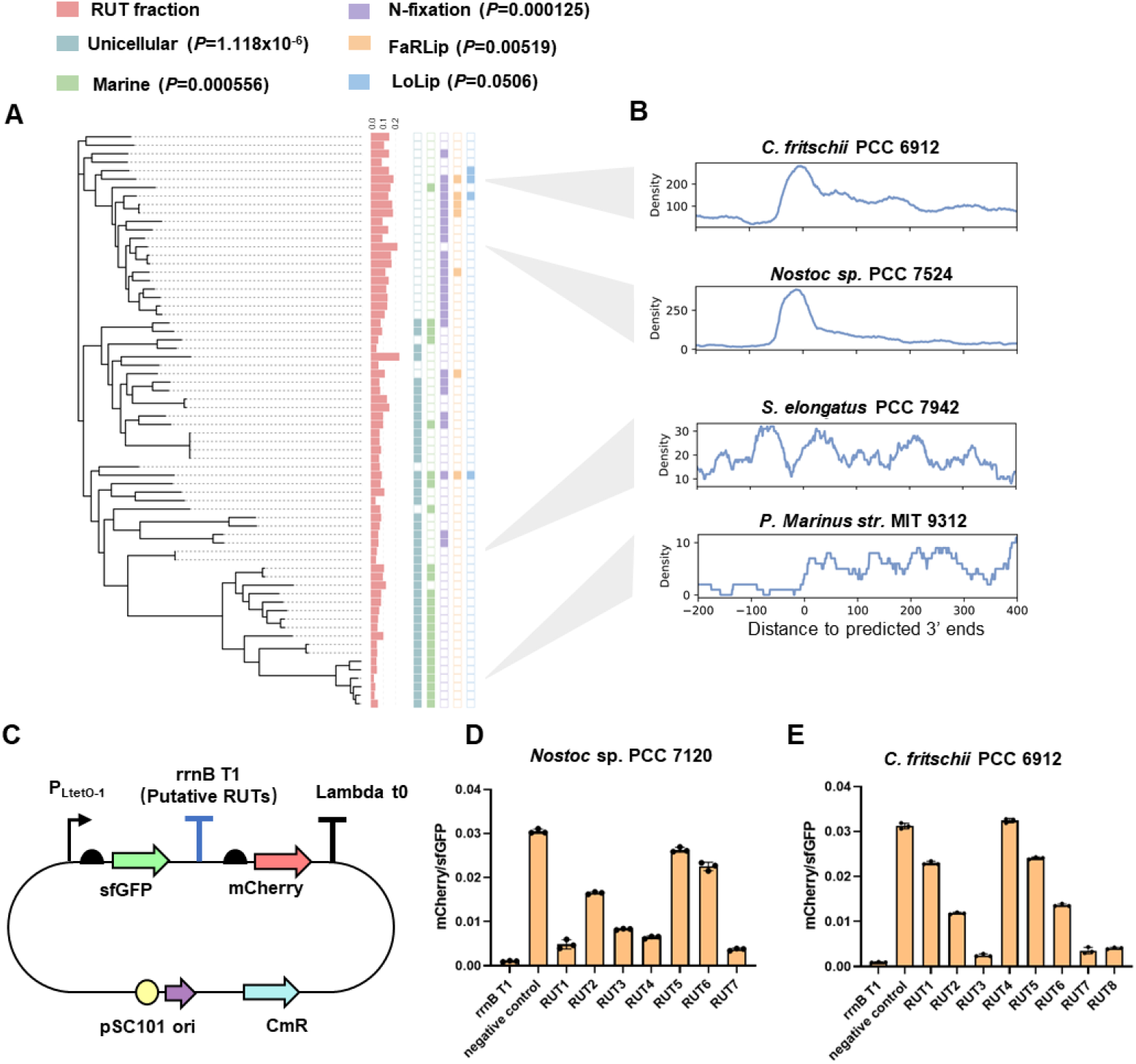
The presence of RUT-like sequences correlates with recently evolved traits in *Cyanobacteria* A. Association between Rho dependency scores and *Cyanobacteria* traits. Wilcoxon test *P*-values were shown in the brackets. B. Metagene plot for predicted RUT sites near predicted transcript 3’ ends at FPR=0.2/KB. Four representative *Cyanobacteria* species are shown. C. Diagram of the reporter plasmid for fluorescence assay. D. mCherry/sfGFP ratio in tested *Nostoc sp*. PCC 7120 RUT sites-like sequences. E. mCherry/sfGFP ratio in tested *C. fritschii* PCC 6912 RUT sites-like sequences. In D and E, error bars show the mean ± SEM of 3 biological replicates.

As *Cyanobacteria* genomes lack functional Rho homologs, here we used a fluorescent reporter assay to test the functionality of these RUT-like sequences in *E. coli* (Figure 5C). In a sfGFP-rrnB T1 terminator-mCherry construct, the rrnB T1 terminator sequence was replaced with either predicted RUT-like sequences or a non-terminating sequence from the lacI gene as a negative control. A low mCherry/sfGFP ratio indicates high transcription termination efficiency. 7 predicted RUT site-like sequences associated with nitrogen fixation-related genes in *Nostoc sp*. PCC 7120 and 8 predicted RUT site-like sequences associated with FarLip and LoLip-related genes in *C. fritschii* PCC 6912 were tested. Among these tested sequences, 7/7 *Nostoc sp*. PCC 7120 sequences and 7/8 *C. fritschii* PCC 6912 sequences have lower mCherry/sfGFP ratios compared to the negative control (Figure 5D and 5E), indicating the predicted RUT sequences are at least partially functional for terminating transcription in *E. coli*.

### Application on identification of AMR genes related premature terminations

As another application of BATTER, we systematically predict premature termination events associated with 5’ UTRs of antimicrobial resistance (AMR) genes. Previous studies have suggested premature transcription termination plays an important role in bacterial transcriptional regulation under antibiotic stress ^[11]^. Premature terminations in bacteria could act in *cis* (through attenuation mechanism) or in *trans* (premature transcription product serve as sRNA to regulate target genes^[18]^ or sponge other sRNAs^[30]^), and in some cases, both mechanisms are simultaneously involved^[47]^. For *cis*-regulation, BATTER-TPE could predict RIT-based premature termination, BATTER-RUT could provide clues for RDT-based premature termination. For *trans*-regulation, as the transcriptional products are functional, their 3’ ends are likely to contain protective stem loops, and could be effectively detected by BATTER-TPE. As a result, although BATTER is designed for predicting primary transcription termination, it can also predict premature termination in many cases. We thus predicted AMR genes in GEMs representative bacterial genomes AMRfinder^[40]^ and identified the premature termination sites in their upstream sequences with BATTER-TPE. The prevalence of predicted upstream terminator structures varied substantially across different AMR gene families, being notably frequent in ribosomal protection proteins, 23S rRNA methyltransferases, and efflux transporters, while rare in other AMR families (Figure 6A-B). It was reported the RppA gene (annotated to ABC-F type ribosomal protection protein by AMRfinder) in *Bacillus thuringiensis* BMB171 is regulated by an attenuator-induced premature termination event^[48]^, our analysis suggests similar regulation is conserved in other ribosome protection protein families. It was well characterized that the ermC gene (annotated to 23S rRNA methyltransferase Erm(C) family by AMRfinder) is regulated by a attenuator that shelters ribosomal binding sites for translational regulation^[49]^, here our predictions suggest in other 23S rRNA methyltransferase families, such as Cfr family 23S rRNA methyltransferase and 23S ribosomal RNA methyltransferase Erm, premature termination-based regulation can be more pervasive. In agreement with our prediction, a previous study also suggested premature terminations of multiple antibiotic efflux-related genes are suppressed upon antibiotic treatment^[11]^. An Rfam search suggested the upstream sequence of macrolide efflux MFS transporter Mef(A) associated premature terminator hits to Enterococcus sRNA_1300 (RF02862), a known sRNA related to antibiotic response^[22]^, suggesting premature termination of these genes may generate *trans*-acting sRNA.

**Figure 6.**
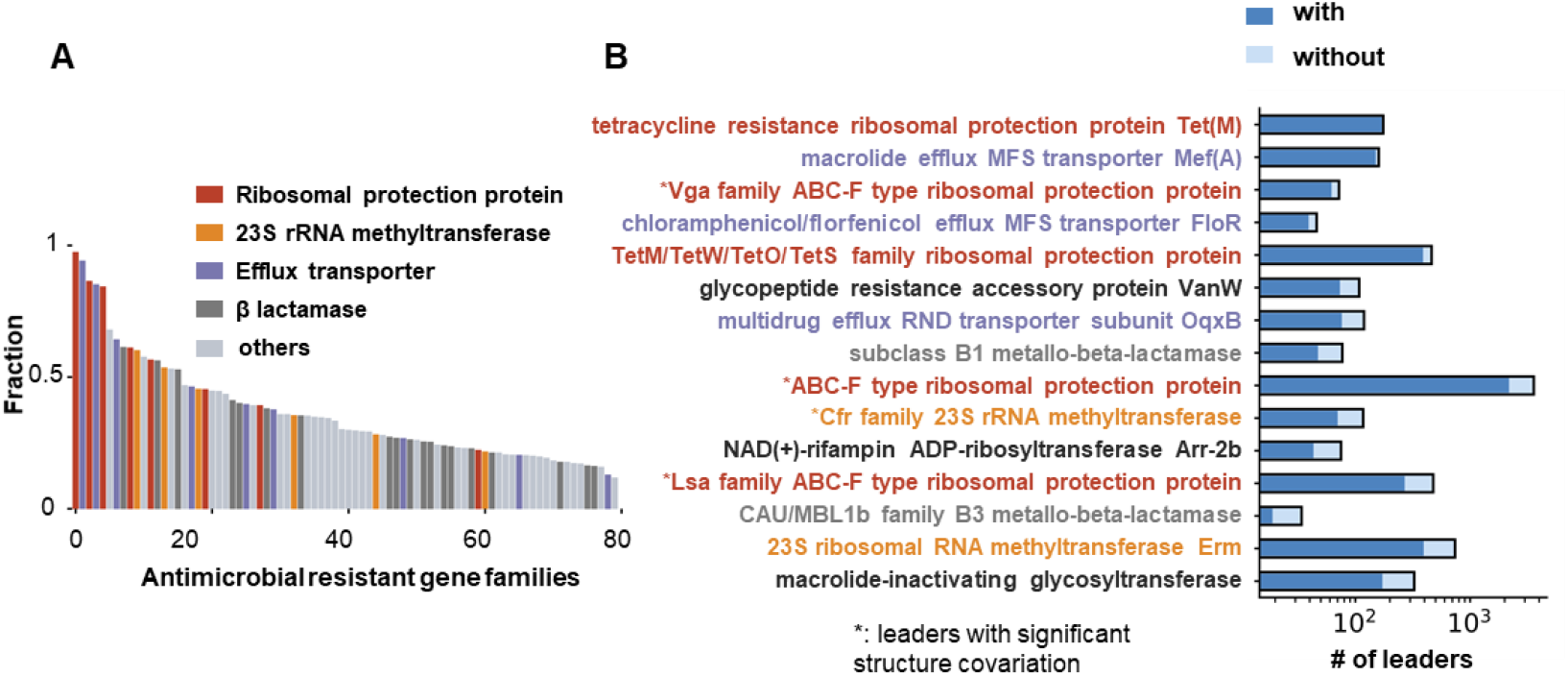
Terminator prediction reveal pervasive riboregulations in some antibiotic resistance gene families A. Antimicrobial resistance (AMR) gene families are ranked by fractions with predicted upstream terminators, each bar indicates an AMR gene family. Gene families that share similar mechanisms of action are labeled with the same color. B. Fraction of leader sequence preceding AMR gene families that are frequently associated with predicted transcription terminators.

For the top 15 AMR gene families with the most predicted upstream terminators, we evaluated the conservation of their secondary structures. Sequence covariation analysis supported conserved secondary structures in upstream sequences of translation protection-related AMRs (ribosomal protection proteins and 23S rRNA methyltransferase), but not those related to efflux transporters (Figure 6B and Figure S8). A potential explanation is that most translation-related premature terminations detected here may act in *cis*, in which RNA structures play a more prominent role; while premature terminations of efflux transporters may generate *trans*-acting sRNAs, which are relatively unstructured.

## Discussion

In this study, by term-seq data analysis, we confirmed that a conserved stem loop structure is often associated with transcript 3’ ends generated by primary termination events in diverse bacterial species, regardless of the dependency of Rho factor. Based on this finding, we developed a data augmentation pipeline and trained BATTER-TPE, a BERT-CRF model for predicting transcript 3’ ends. We also developed an accessory module, BATTER-RUT, for detecting putative RUT sites. Benchmarking BATTER on diverse datasets demonstrated its advantage compared to existing tools. Using BATTER, we systematically characterized the properties of 3’ ends associated stem loops and evaluated the role of Rho-dependent termination across major bacterial lineages. Additionally, we systematically predicted putative transcript 3’ ends located upstream of AMR genes, identifying several AMR gene families potentially regulated by premature termination.

While BATTER-TPE outperforms existing tools, it does have some limitations. BATTER-TPE is optimized to detect 3’ ends from transcription termination events in which the resulting products are functional and hence should be protected from RNase degradation, as in the case of primary transcription termination of protein-coding genes and functional ncRNAs. Many non-primary termination events, however, do not follow this paradigm. In addition, archaea terminators usually do not contain stem loop structures^[50]^, and BATTER-TPE cannot effectively predict transcript 3’ ends in archaea. Thirdly, although BATTER-RUT mildly outperformed existing tools for RUT prediction and achieved reasonable performance for detecting 3’ end-associated RUTs in combination with BATTER-TPE, exhaustive genome-wide prediction of RUTs remains challenging, and comparative analysis of Rho protein function other than terminating primary transcription is also an interesting direction for future investigation.

Our systematic prediction of 3’ end-associated stem loops revealed that inter-species variations generally exceed inter-genes variations. The RNA polymerase complex, while universally present in bacteria with conserved features such as subunit composition, has evolved clade-specific variations that affect transcription regulation^[51]^. For instance, β and β’ subunits of RNA polymerase contain clade-specific insertions; deletions of these regions lead to various functional deficiencies^[52]^. Multiple clade-specific transcription factors have also been reported to regulate bacterial transcription^[51]^. Assuming sequences and structures of 3’ end-associated stem loops are optimized for stalling the RNA polymerase complex, inter-species variations of terminators may reflect co-evolutions between terminators and the transcription machinery.

The discovery of RUT-like signatures near transcript 3’ ends in certain *Cyanobacteria* species is very intriguing. One potential explanation is that those *Cyanobacteria* species may harbor an undetected Rho homolog. It has been reported that in *Arabidopsis thaliana*, RHON1, a Rho-like protein, is responsible for some transcription termination events in plastids^[53]^. It was well accepted that plastids in plants evolve from endosymbiotic *Cyanobacteria*^[54]^; hence, some *Cyanobacteria* may harbor a Rho-like protein homolog. Following a previously proposed strategy^[35]^, in this study, proteins with both Rho RNA binding domain and ATPase domains are considered as functional Rho, and no functional Rho was detected in *Cyanobacteria* clade. We could not completely exclude the probability that some *Cyanobacteria* species may harbor homologs of Rho protein, as if the sequence divergence is too large, Rho protein homologs may elude detection. An alternative hypothesis is that Rho protein was lost in the common ancestor of existing *Cyanobacteria*, in some descendants, other protein factors may have adapted to use the original Rho binding sites for transcription termination, while in others, RUT sites may have been lost entirely. This may explain the association found between RUT-like signatures and recently evolved traits, such as nitrogen fixation, FarLip, and LoLip. Both hypotheses require further experimental validation.

BATTER-TPE is optimized for predicting 3’ ends of the primary transcripts. We have shown in practice it can also be applied to predict a considerable fraction of premature termination events, including termination events that generate functional products (such as *trans*-acting sRNAs generated by both RITs and RDTs), and *cis*-regulatory RITs. Although BATTER-RUT could detect some RUT sites in leader sequences, comprehensive prediction of all RUTs across the bacterial genome remains an open problem. Despite such limitations, BATTER successfully detects variants of known and novel riboregulators. These riboregulators can serve as potential drug targets for antibiotic-resistant strains. Besides premature termination events, at downstream sequences of some protein-coding genes, multiple terminators exist; terminators also exist in some intra-operonic regions. The selection between these alternative terminators indicates regulatory events, is an interesting direction for further investigation.

As BATTER is optimized for 3’ ends of functional transcripts in very diverse bacteria species, taking genome sequences as the solitary input, it is particularly useful for discovering 3’ ends of functional transcripts in contigs assembled from metagenome samples, where very diverse bacteria lineages, including uncultivated ones, are often present. We hope BATTER will be a useful tool for understanding the diversity of ncRNAs across the bacterial domain.

## Code availability

Codes of BATTER, data for model training, scripts for model benchmarking, and predicted transcript 3’ ends in GEMs genomes are available in zenodo (https://zenodo.org/records/16761763).

## Acknowledgments

This work is supported by National Key RD Program of China (2022YFA1304200), National Natural Science Foundation of China (32170671,8237061277), Tsinghua University Guoqiang Institute Grant (2021GQG1020), Tsinghua University Initiative Scientific Research Program of Precision Medicine (2022ZLA003), Bioinformatics Platform of National Center for Protein Sciences (Beijing) (2021-NCPSB-005). This study was also supported by Bayer Micro-funding, Beijing Advanced Innovation Center for Structural Biology, Bio-Computing Platform of Tsinghua University Branch of China National Center for Protein Sciences.

## Notes

**Competing Interest Statement:** The authors declare no potential conflicts of interest.

https://zenodo.org/records/16675150

### Competing Interest Statement

The authors have declared no competing interest.

### Summary of Updates

(1) Revise the rationales for defining primary TESs. (2) Update data availability.

