## Supplementary information for "Conserved 3’ Stem-Loop Structures Enable Comprehensive Analysis of Bacterial Transcription Termination in Metagenomes"

### Conserved 3’ Stem-Loop Structures Enable Comprehensive Analysis of Bacterial Transcription Termination in Metagenomes, Regardless of Rho Factor Dependency

Yunfan Jin, Jiyun Cui, Hongli Ma, Fei Gan, Zhenjiang Zech Xu, Zhi John Lu

Zhi John Lu

Zhenjiang Zech Xu

Fei Gan


### Supporting Information Text

#### The data augmentation pipeline

The data augmentation pipeline is based on three assumptions: 1) the vast majority of transcript 3' ends are associated with hairpin structures; 2) a single covariance model[1] is sufficient to model the variations between terminators of homologous genes in closely related species; 3) in species lacking experimental data, genes homologous to those associated with experimentally supported 3' ends are more likely to associated with terminators compared to random protein-coding genes.

Term-seq defined 3' ends located within the downstream 128 nt of a protein-coding gene on the same strand were annotated as primary 3' ends. As shown in Figure S2 A, for the collected 3' end mapping data, protein products of genes with primary 3' ends assigned were clustered to 40% sequence identity with cd-hit[2]. 115 clusters that contain protein sequences from at least 3 bacterial phyla were annotated with hmmsearch[3] (hmmer version 3.3.2) using Pfam[4] database and default parameters (Dataset S1). For homolog search, we used Genomic catalog of Earth's Microbiomes (GEMs), a diverse prokaryotic genome collection, which clustered metagenome-assembled genomes (MAGs) and refseq genomes into 45,599 species-level representatives, of which 42,905 are bacterial species[5]. Pfam hits of different protein clusters were utilized as surrogates for homology search. If a protein sequence was annotated to at least 3/4 Pfam domains that were assigned to a gene cluster, it was mapped to the cluster for downstream analysis.

For protein hits, sequences of 16 nt upstream and 128 nt downstream of the stop codons were extracted and subjected to CMfinder[6] search. As we focused on primary termination events, the 128 nt cutoff for the downstream sequence was chosen. 16 nt upstream sequences were padded, aimed to cope with the scenarios where the stem loop structures occasionally overlap with protein-coding genes. To consider inter-species variation and reduce the computational burden, candidate sequences of each cluster were further grouped by the taxonomy relationship. Phyla with < 16 sequences were neglected; phyla with 16-128 sequences were grouped as CMfinder[6] input; for phyla with > 128 sequences, we further grouped these sequences by lower taxonomy level (orders), and ran the previous process recursively (from phylum, class, order, family, genus, to species). This process guarantees each CMfinder run receives no more than 128 sequences from phylogenetically related species.

To ensure reliable data augmentation, the cm models were filtered using the following criteria:

(1) In addition to 3' end-associated stem loops, CMfinder can detect other conserved structural elements, hence only covariance models with exactly 1 stem-loop, and at least 5 base pairs were reserved.

(2) Based on the assumption that stem loops conserved across multiple gene families are more reliable, all candidate downstream sequences were searched by covariance models that passed the first criterion with cmsearch program in infernal[1] (version 1.0.2). Only cm models that hit more than 40 downstream sequences in at least 15 other protein clusters were kept.

To augment terminator instances with ncRNAs, Rfam seed alignments of a list of known bacteria ncRNA (mainly attenuators and sRNAs, Dataset S1) were searched against GEMs bacteria genomes with mmseqs[7], and cmsearch[1] was used to refine the alignment boundary. The resulting sequences, together with their downstream 128 nt sequences, were searched using cmsearch with cm models passed filtering, and a similar process was applied to downstream sequences of a random subset (128 genes per genome) of protein-coding genes to identify putative terminators of diverse protein-coding genes.

The secondary structures of all putative terminators were predicted with RNAfold[8], and further filtered by the number of base pairs (>7), and the number of loops (exactly 1). The stem-loops with 5 nt flanking sequences at both sides were defined as terminators for model training.

#### Implementation and training and BATTER-TPE

The longformer-CRF model was implemented using pytorch[9], with longformer encoder (8 layers with hidden size of 256, intermediate size of 1024, max length of 514, and 64 nt sliding window attention at each side) implemented in transformers[10] package. The model was pre-trained on the 42,905 GEMs representative genomes with masked language modeling (MLM) loss[11], that is, 20% of the tokens are masked, and the model is trained to predict the masked tokens with unmasked tokens (the loss is defined as cross entropy between predicted logits and actual tokens).

Denoting longformer predicted score for state j (0 for background, 1 for terminator) at position $i$ as $E_{ij}$, state of position $i$ as $T_i$. The learnable parameters in CRF layers are transition scores $P_{T_{i-1}T_i}$ between two token states. Both longformer predicted scores and transition scores can be interpreted as probabilities in log space. At the training stage, the state of each token is known, and model parameters are trained end-to-end, the object is to maximize the likelihood of the given state configurations. The score of given state assignments $T = (T_1, \ldots, T_N)$ can be written as:

$$score_T = \sum_{i=1}^{N} P_{T_{i-1}T_i} + E_{iT_i}$$

The likelihood of this state assignment is:

$$LL_T = \frac{\exp\,(score_T)}{\sum_{t \in all\ states} \exp(score_t)}$$

Maximizing the likelihood is equivalent to minimizing negative log-likelihood, which is the loss function:

$$loss = score_T - \log\,(\sum_{t \in all\ states} \exp(score_t))$$

For both model pretraining and training, the AdamW optimizer (lr:5e-5, eps:1e-8, betas:(0.9,0.999) ) was used with a weight decay of 0.01 and linear decayed learning rate scheduling. A batch size of 128 was used. For model training, 70% of the instances contain putative terminators, and the remaining 30% were either random genomic sequences (25%) or random genomic sequences shuffled by 4-mer with ushuffle[12] (75%). For each instance, a subsequence with length uniformly distributed between 64nt and 510 nt was cropped as model input.

In the inference stage, state paths with top k $score_T$ were determined by Viterbi decoding. Spans of consecutive tokens that start from position $s$ and end at position $e$ were tagged as a terminator that is longer than 32 nt were identified, and averaged predicted probabilities of the tokens were utilized to reflect the confidence $C(s..e)$ of the prediction:

$$C(s..e) = \frac{1}{e-s+1}\sum_{i=s}^{e}\frac{\exp\,(E_{i1})}{\exp(E_{i0}) + \exp\,(E_{i1})}$$

Input genome sequences were scanned with a window size of 500 nt, and a stride size of 100 nt. The top 10 predictions were considered. For overlapping predictions on the same strand, only the best-scoring one is selected as the final prediction.

#### Implementation of BATTER-RUT

As shown in Figure S3, to detect putative Rho binding sites, the input sequences were scanned with a window size of 100 nt and a stride length of 40 nt. All YC dimers were extracted from the 100 nt sequence. Among these YC dimers, some are embedded in left-side flanking sequences of RUT sites (denoted as "L"), some are embedded in right-side flanking sequences of RUT sites (denoted as "R"). Within the RUT site, some YC dimers are the relatively conserved 3' ends in the substrate of Rho monomer (denoted as "C"), and the remaining are not conserved (denoted as "N"). As shown in Figure S3 A, when scanning from left to right, feasible transitions include L to L, L to C, C to N, N to N, N to C, C to C, C to R, and R to R. To identify the subset YCs that is most likely to correspond to "C" state, we defined a scoring rule, given states assignment of all YCs, neighboring Cs are scored according to their distance $d$ (Figure S3 B). The scoring rule is defined as:

$$S(d) = -0.25e^{-\frac{d}{10}} + \sum_{i=-1}^{1} N(d-10; i, 0.5) + e^{-1}\sum_{i=-1}^{1} N(d-20; i, 0.5)$$

When setting the maximum score to 1, the normalized score become:

$$NS(d) = S(d)/S(10)$$

Here $N(x;\mu,\sigma^2)$ indicates Gaussian with mean of $\mu$ and variance of $\sigma^2$. Normalized scores of all neighbors are sum up to produce final score. A distance greater than 25 nt produces a large penalty. This scoring rule encourages selected adjacent YC dimers to have distances close to 10nt or 20nt (to tolerate scenarios when the substrate of Rho monomer does not end with YC). The state assignments that maximize the scoring rule were identified using a beam search heuristic: all YC dimers were scanned from left to right, each YC was assigned as either "C" or one of "L", "R", and "N". Only ttransitionsbetween two "C" states contribute to the scoring. We only kept track of the top k (set to 500 by default) state assignments, for the best assignments at the last position, positions of YC dimers assigned to "C" states were selected as candidate RUT sites. Candidate RUT sites with at least 4 properly spaced YC dimers were subsequently rescored by $\frac{\#YC}{L} + \log_2 C/G$, where $\#YC$ is the number of YC dimers, $L$ is the length of the candidate sequence. Candidates with $C/G < 1$ were not considered (Figure S3 C). For sequence flanking 3' ends predicted by BATTER-TPE, BATTER-RUT could detect whether there are RUT site-like sequences.

#### Performance evaluation

For performance evaluation, the false positive rate was estimated by scanning 1000 sequences (1KB in length) sampled from the tetramer frequency distribution of the corresponding genomes.

Predicted intervals that overlap with the upstream 40 nt of term-seq identified 3' ends on the same strand were considered as true positives.

For BATTER-TPE and BATTER-RUT, the default settings were used. For RNIE, the parameters "--thresh 0 --gene -m gene.cm" were used. For transterm-HP, the parameters "--min-conf=50 --all-context" were used. For RhoTermPredict, the default parameters were used. For the OPLS-DA model, as the original publication does not provide model weights, the 111 features were extracted using the MakeDescriptors.py script[13], and an OPLS-DA model was retrained using validated Rho sequences used in the original study as positive instances, and dinucleotide shuffled sequences from the same genome as negative instances. The genome sequences were scanned with a stride length of 50 nt, and a window size of 500nt for OPLS-DA predictions. For all tools considered, the intervals with the highest score in overlapped predictions (if any) were selected for performance evaluation.

The data augmentation pipeline starts from 115 gene clusters with term-seq supports spanning multiple phyla, to avoid over-estimation of model performance potentially introduced by bias toward these genes, for all species, 3' ends of genes with significant hits to these genes clusters by mmseqs[7] search with alignment coverage > 50% were also excluded in performance evaluation. Although *E. coli* and *B. subtilis* data are present in the training set, as only 3' ends associated with the 115 gene clusters were used for data augmentation, and model training only used the augmented data, there was no information leakage. For *B. burgdorferi* and *M. tuberculosis*, all 3' ends were never seen during model building.

#### Analysis of variations of predicted stem loops

The prediction performance and terminator characteristics were visualized with interactive Tree of Life [14] (https://itol.embl.de/). Predicted secondary structures of representative terminators were visualized with varna[15].

For t-SNE visualization of terminators, a BERT encoder was trained to project each RNA sequence to a numeric vector on a 256-dimensional unit sphere. The model was implemented with transformers[10] library, pre-trained on RNAcentral[16] database with MLM loss, and fine-tuned on Rfam supervised by a triplet loss. Given an instance in a minibatch as anchor $a$, within the same minibatch, the instance that shares the same label with the anchor while having the largest cosine distance $d_{pa}$ is considered as the positive instance $p$, and the instance with the different label while having the smallest cosine distance $d_{na}$ is considered as the negative instance $n$ (Figure S7 A). Then triplet loss of the anchor instance is defined as:

$$L(p, n, a) = max\ (d_{pa} - d_{na} + margin, 0)$$

The hyperparameter $margin$ is set to 0.32. The qualities of RNA embeddings were validated on holdout RNA families. The vector embeddings were visualized with Multicore-TSNE[17].

PCA and K-means clustering were implemented with decomposition.PCA and cluster.KMeans modules in scikit-learn package[18], respectively. The statsmodels[19] package was used for the linear model that decomposes the variance of the principal components.

### Supplementary Figures

#### Figure S1.

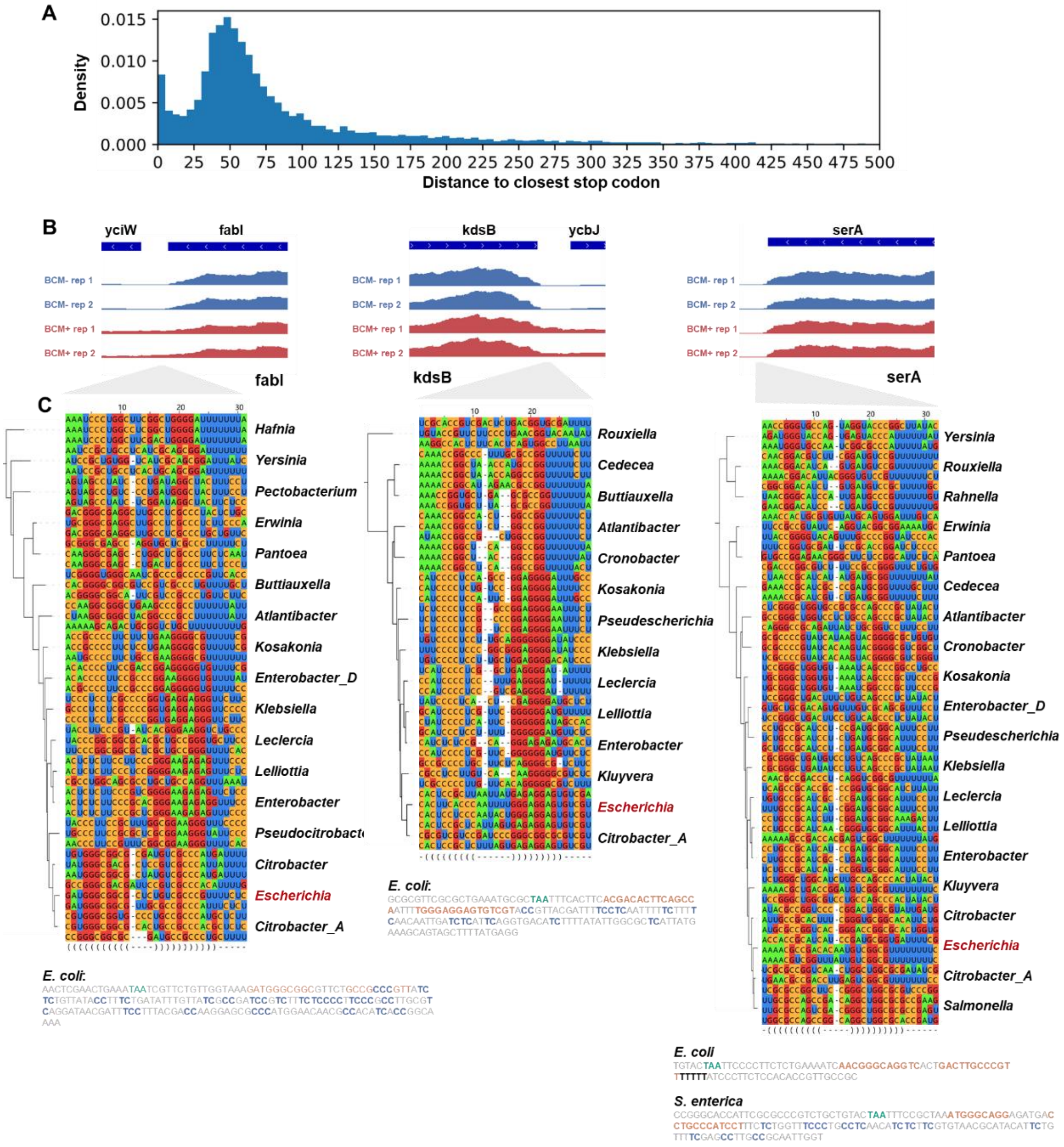

A. The distribution of distances between TESs and their closest upstream stop codons.

B. The read coverages of the BCM-treated samples and control samples are shown. According to RNA-seq data, in *E. coli*, the transcription termination of fabI gene and kdsB gene depends on the Rho factor, while the transcription termination of serA gene is Rho-independent.

C. Multiple sequence alignments of conserved stem loops downstream fabI, kdsB, and serA genes in *Enterobacteriaceae* clade are shown. In each of the 3 genes, downstream sequences of the stem loops in some species have RIT-like properties, while the others have RDT-like properties.

**Figure S2.**

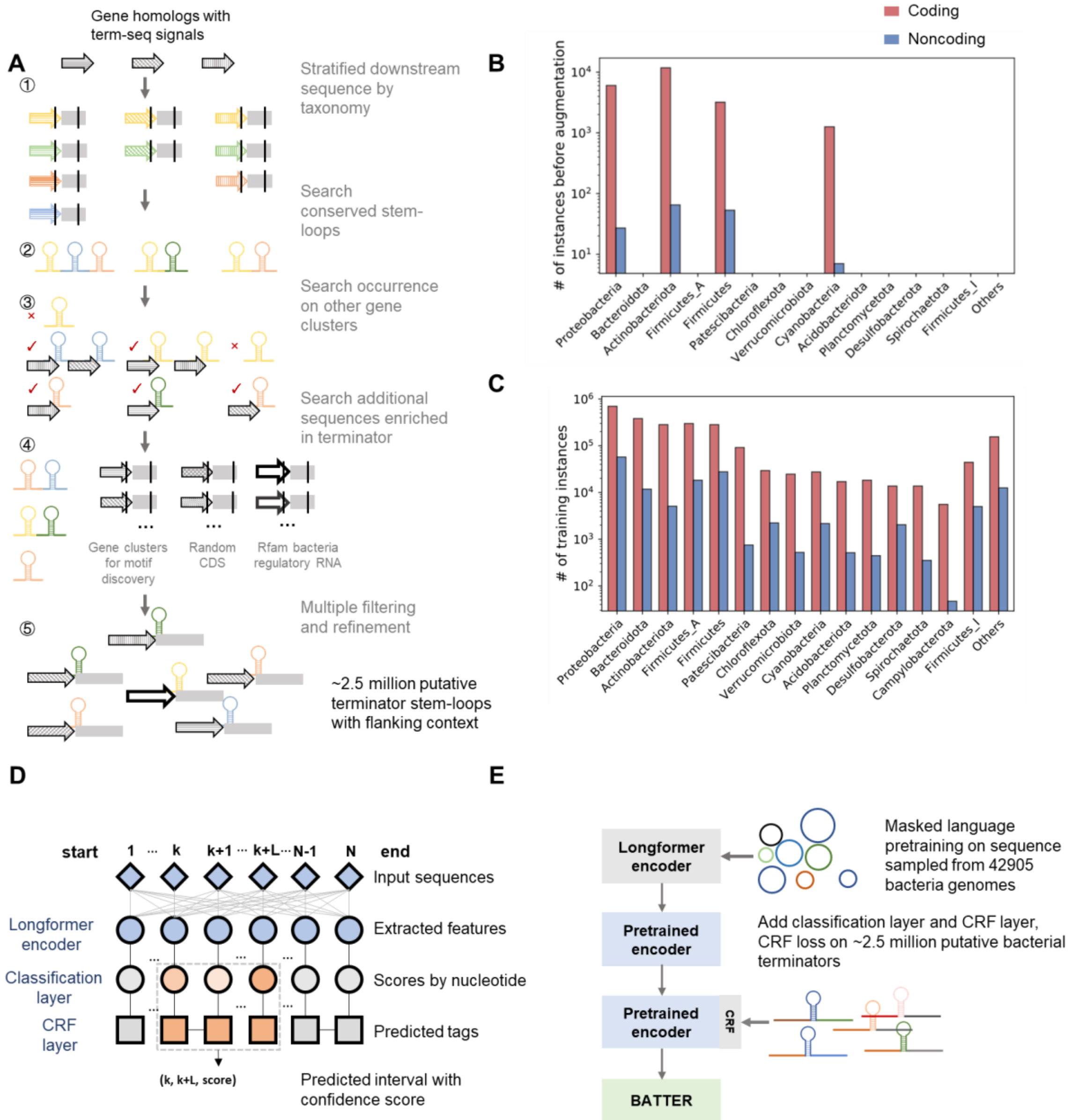

A. A schematic overview of the data augmentation pipeline. For each protein-coding gene cluster (patterns indicate different gene clusters) supported by term-seq signal from at least 3 phyla, 16 nt upstream and 128 nt downstream sequences relative to stop codons were extracted and stratified by taxonomy (step ①, colors indicate different clades), then subjected to CMfinder search (step ②). Stem-loop motifs discovered from each cluster were searched against other gene clusters, and motifs specific to too few gene clusters were discarded (step ③). Motifs passed the filter were further searched against diverse sequences known to be enriched in terminators (step ④). After multiple steps of filtering and refinement, a final set of putative terminators was identified (step ⑤).

B. Total number of experimentally supported primary 3' ends.

C. Total number of putative 3' ends that are associated with stem loops after data augmentation.

D. The model architecture of BERT-CRF for terminator prediction. The model is composed of a longformer encoder, a dense layer for classification, and a CRF layer. The logits predicted by longformer model are post-processed by a CRF layer to make the final prediction. For predicted intervals, the average of probabilities by nucleotide (predicted by the classification layer) across the interval is utilized as a confidence measure.

E. The model training pipeline. The longformer encoder is pre-trained on bacteria genomic sequences sampled from GEMs bacteria genomes with MLM loss in an unsupervised manner. The encoder parameters, together with parameters of the appended classification layer and CRF layer, are jointly fine-tuned to predict transcription terminators.

**Figure S3.**

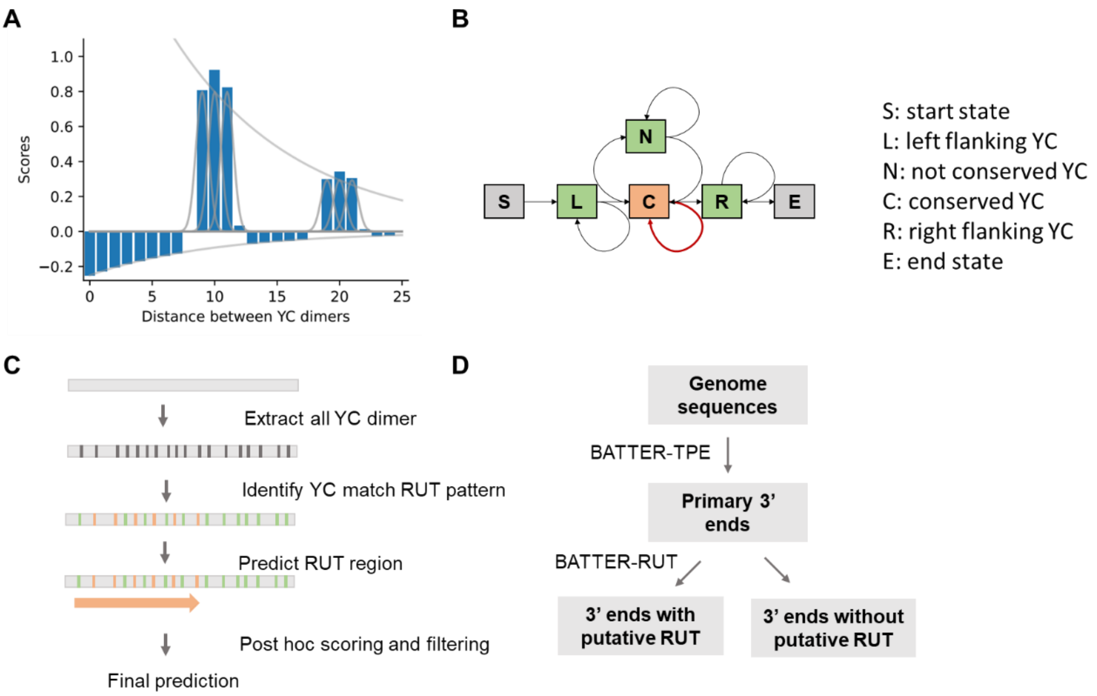

A. Schematic overview of RUT sites prediction algorithm. Subsets of up to 6 YC dimers that maximize the scoring rule were identified using a beam search heuristic, candidate RUT sites with at least 4 properly spaced YC dimers were subsequently rescored.

B. Feasible transitions between the conserved 3' YC dimers (state C) and other non-conserved YC dimers (states L, N, and R).

C. The scoring rule is based on the distance between YC dimers. The scoring rule encourages selected adjacent YC dimers to have distances close to 10 nt, or 20nt (to tolerate scenarios when the substrate of the Rho monomer does not end with YC).

D. BATTER-RUT as a post-processing module for 3' ends predicted by BATTER-TPE.

**Figure S4.**

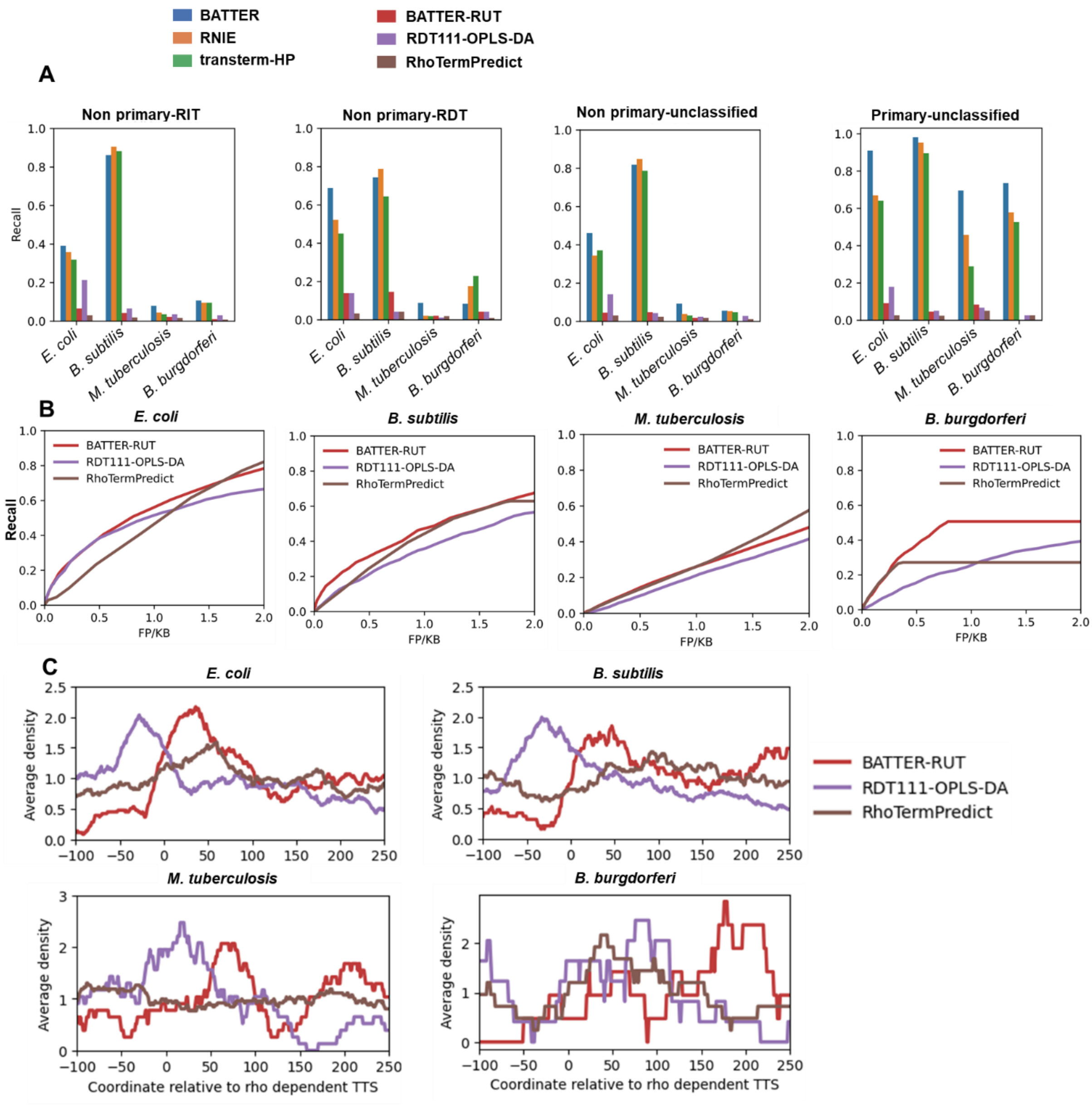

A. Recalls of stem loop prediction-based methods (BATTER-TPE, RNIE, and transterm-HP) and RUT sites prediction-based methods (BATTER-RUT, RDT111-OPLS-DA and RhoTermPredict) at FPR of 0.1/KB, grouped by RIT, RDT, and unclassified. Non-primary 3' ends and unclassified primary 3' ends are shown here.

B. Recalls of 3 RUT sites prediction methods in identifying Rho-dependent regions at different FPR cutoffs in 4 species.

C. Metagene plot for locations of predicted RUT sites near transcripts' primary 3' ends in 4 species.

**Figure S5.**

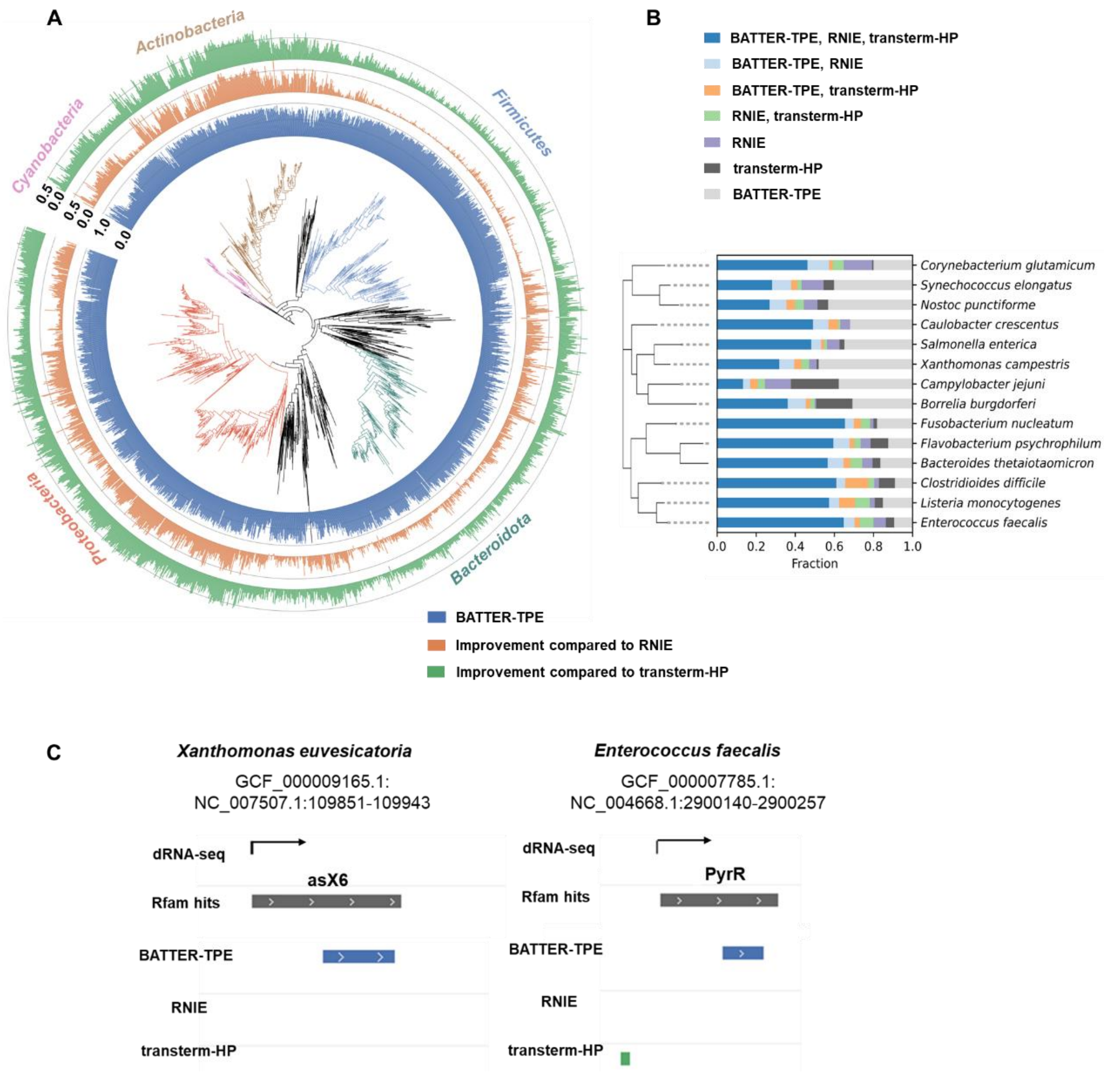

A. Performance comparisons on predicting terminators between tail-to-tail gene pairs in diverse bacterial species. The inner cycle shows recalls of BATTER-TPE (at an FPR of 0.1/KB), and the outer two cycles show improvements in recalls relative to RNIE and transterm-HP, respectively.

B. Overlaps between RNIE, BATTER, and transterm-HP's predictions of transcript 3' ends located downstream of TSSs annotated by curated dRNA-seq datasets.

C. Two TSS-associated terminators that were only predicted by BATTER, which matched Rfam families that were withheld from data augmentation for ncRNA terminators were showcased.

**Figure S6.**

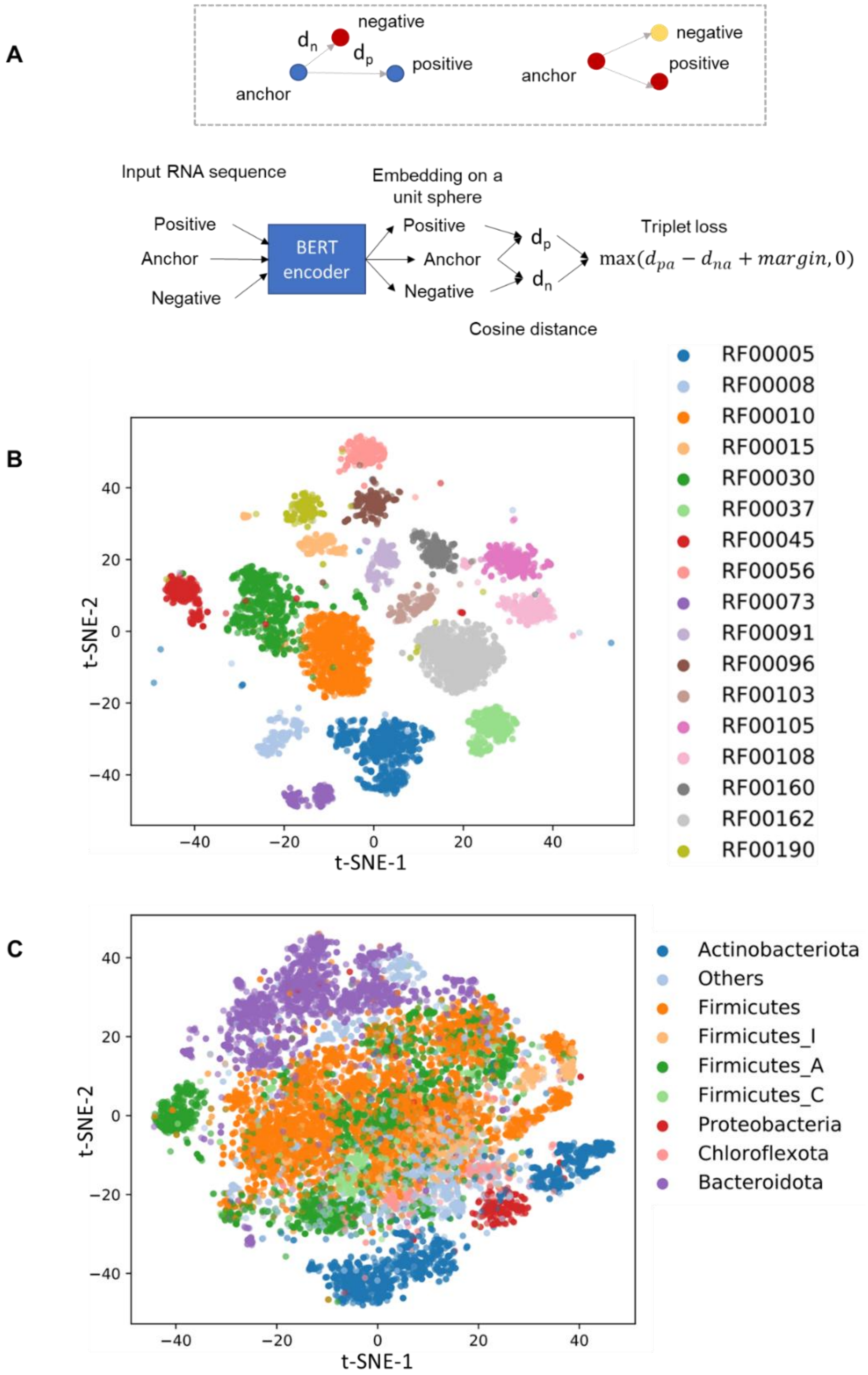

A. Schematic illustration of RNA encoder. A BERT model pretrained on RNAcentral database is finetuned using Rfam data under the supervision of triplet loss that pushes RNA sequences of the same family closer to each other.

B. t-SNE visualization of the embedding of RNA in the holdout set, colors indicate different RNA families.

C. t-SNE visualization of SAM riboswitches (holdout from training), colors indicate different phyla.

**Figure S7**

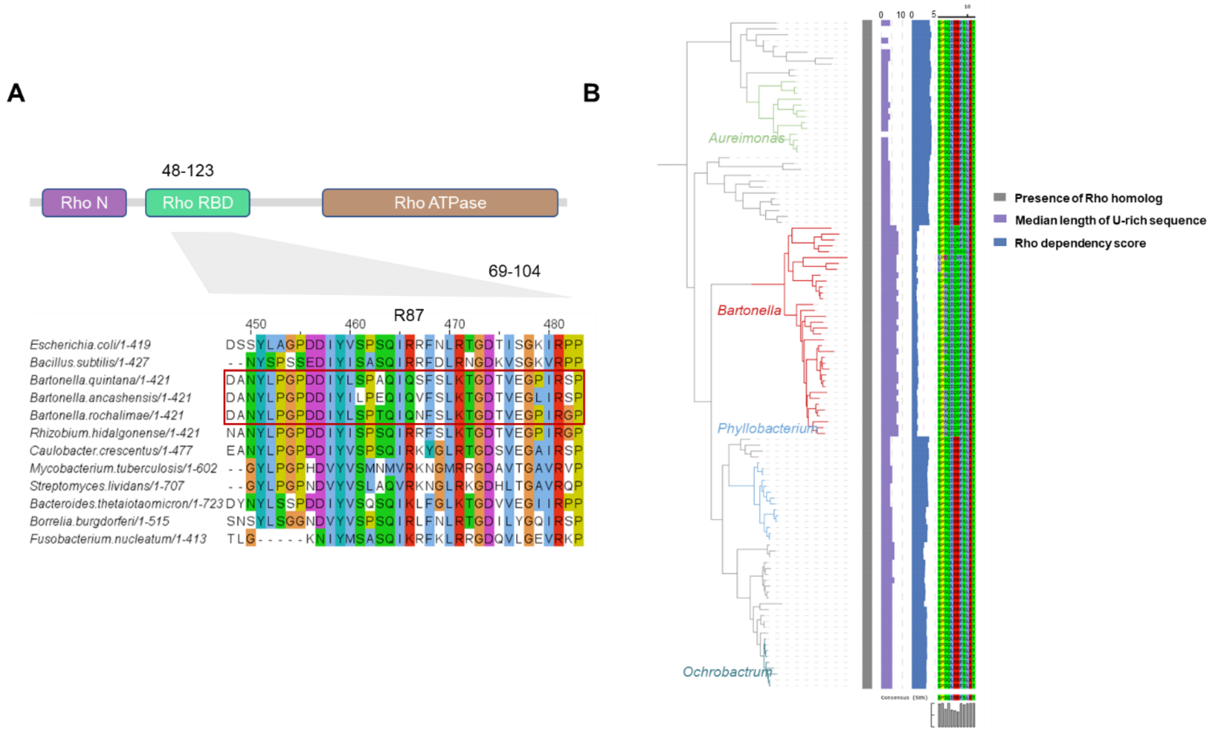

A. The highly conserved basic residue mapped to R87 in *E. coli* Rho protein is substituted into Q in *Bartonella* species.

B. *Bartonella* species have a sharp drop in Rho dependency scores compared to closely related genera in *Rhizobiaceae* family.

**Figure S8**

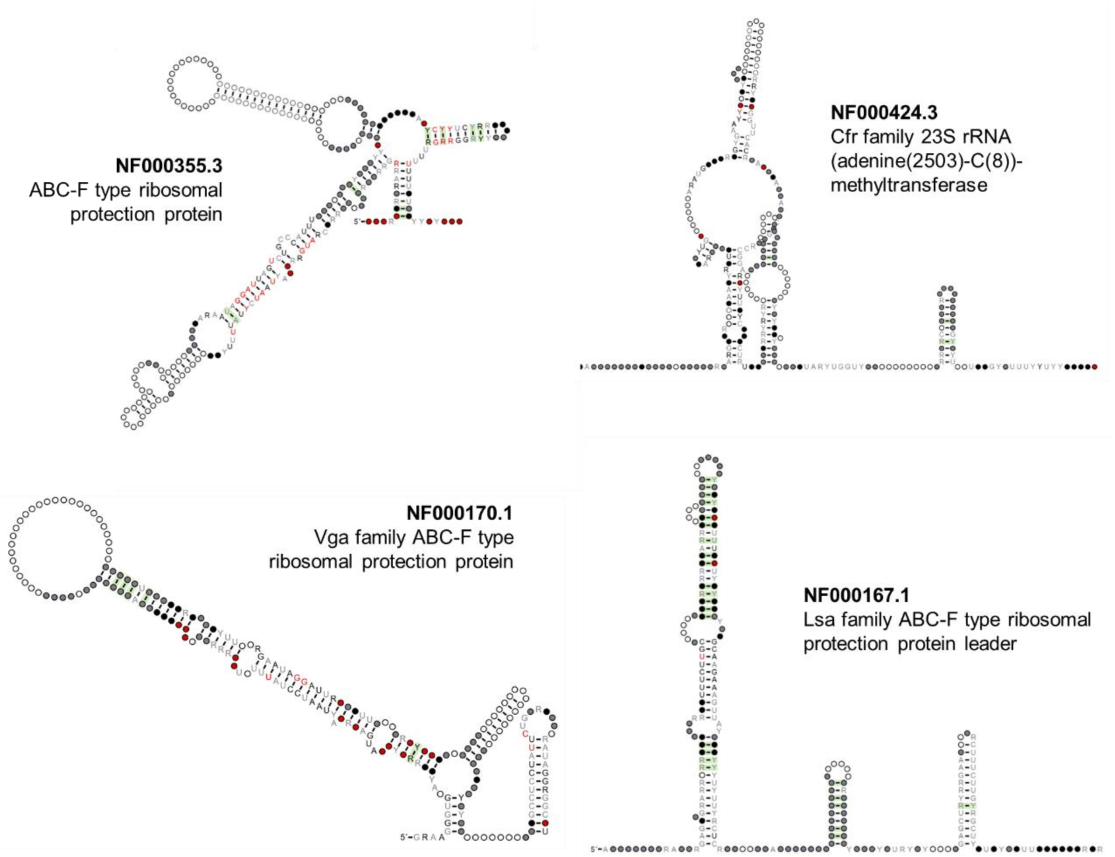

Predicted consensus RNA secondary structures of noncoding RNAs with significant structural covariations upstream of 4 AMR gene families.

### Supplementary Tables

#### Table S1

**Curated 3' ends mapping data**

| Published year | Species | Phylum | Reference genome | PMID | Used for data augmentation |
|---|---|---|---|---|---|
| 2018 | *Escherichia coli str. K-12 substr. MG1655* | *Proteobacteria* | GCF_000005845.1 | 29606352 | TRUE |
| 2018 | *Escherichia coli str. K-12 substr. MG1655* | *Proteobacteria* | GCF_000005845.2 | 38030608 | FALSE |
| 2018 | *Bacillus subtilis subsp. subtilis str. 168* | *Firmicutes* | GCF_000009045.1 | 29606352 | TRUE |
| 2018 | *Caulobacter vibrioides NA1000* | *Proteobacteria* | GCF_000022005.1 | 29606352 | TRUE |
| 2018 | *Vibrio natriegens NBRC 15636* | *Proteobacteria* | GCF_001456255.1 | 29606352 | TRUE |
| 2018 | *Streptococcus pneumoniae TIGR4* | *Firmicutes* | GCF_000006885.1 | 30517198 | TRUE |
| 2019 | *Streptomyces lividans TK24* | *Actinobacteria* | GCF_000739105.1 | 31555254 | TRUE |
| 2019 | *Pseudomonas aeruginosa PAO1* | *Proteobacteria* | GCF_000006765.1 | 31594819 | TRUE |
| 2020 | *Zymomonas mobilis subsp. mobilis ZM4* | *Proteobacteria* | GCF_003054575.1 | 32694125 | TRUE |
| 2020 | *Streptomyces avermitilis MA-4680* | *Actinobacteria* | GCF_000009765.2 | 33319794 | TRUE |
| 2020 | *Streptomyces griseus subsp. griseus NBRC 13350* | *Actinobacteria* | GCF_000010605.1 | 33319794 | TRUE |
| 2020 | *Streptomyces coelicolor A3(2)* | *Actinobacteria* | GCF_000203835.1 | 33319794 | TRUE |
| 2020 | *Streptomyces lividans TK24* | *Actinobacteria* | GCF_000739105.1 | 33319794 | TRUE |
| 2020 | *Streptomyces tsukubensis* | *Actinobacteria* | GCF_003932715.1 | 33319794 | TRUE |
| 2020 | *Streptomyces clavuligerus* | *Actinobacteria* | GCF_005519465.1 | 33319794 | TRUE |
| 2020 | *Streptomyces venezuelae* | *Actinobacteria* | GCF_015710995.1 | 33319794 | TRUE |
| 2021 | *Streptomyces clavuligerus* | *Actinobacteria* | GCF_005519465.1 | 33947798 | TRUE |
| 2021 | *Synechocystis sp. PCC 7338* | *Cyanobacteria* | GCF_018282115.1 | 34054774 | TRUE |
| 2022 | *Synechocystis sp. PCC 6803* | *Cyanobacteria* | GCF_000009725.1 | 34874777 | TRUE |
| 2022 | *Dickeya dadantii 3937* | *Proteobacteria* | GCF_000147055.1 | 35491820 | TRUE |
| 2023 | *Borreliella burgdorferi B31* | *Spirochaetota* | GCF_000008685.2 | 37402717 | FALSE |
| 2023 | *Mycobacterium tuberculosis H37Rv* | *Actinobacteria* | GCF_000195955.2 | 37096044 | FALSE |

**Table S2**

**Curated RNA-seq data with Rho inhibition**

| Published year | Species | GC content | Phylum | Reference genome | Accession | PMID |
|---|---|---|---|---|---|---|
| 2012 | *Escherichia coli str. K-12 substr. MG1655* | 51% | *Proteobacteria* | GCF_000005845.2 | GSE41939 | 23207917 |
| 2023 | *Borreliella burgdorferi B31* | 28% | *Spirochaetota* | GCF_000008685.2 | GSE222085 | 37402717 |
| 2023 | *Bacillus subtilis subsp. subtilis str. 168* | 43.50% | *Firmicutes* | GCF_000009045.1 | GSE195579 | 36735730 |
| 2023 | *Mycobacterium tuberculosis H37Rv* | 65.50% | *Actinobacteria* | GCF_000195955.2 | E-MTAB-11753 | 37096044 |

#### Table S3

##### Considered species in metaterm-seq dataset

| Phylum | Class | Order | Family | Genus | Species | Reference genome used |
|---|---|---|---|---|---|---|
| Firmicutes | Bacilli | Lactobacillales | Streptococcaceae | Streptococcus | Streptococcus_sanguinis | GCF_000014205.1 |
| Firmicutes | Bacilli | Lactobacillales | Streptococcaceae | Streptococcus | Streptococcus_gordonii | GCF_000017005.1 |
| Actinobacteria | Actinobacteria | Corynebacteriales | Corynebacteriaceae | Corynebacterium | Corynebacterium_matruchotii | GCF_000158635.1 |
| Firmicutes | Bacilli | Bacillales | Bacillales_unclassified | Gemella | Gemella_morbillorum | GCF_000185645.1 |
| Firmicutes | Bacilli | Bacillales | Bacillales_unclassified | Gemella | Gemella_haemolysans | GCF_000204355.1 |
| Firmicutes | Bacilli | Bacillales | Bacillales_unclassified | Gemella | Gemella_sanguinis | GCF_001052115.1 |
| Firmicutes | Negativicutes | Veillonellales | Veillonellaceae | Veillonella | Veillonella_parvula | GCF_016127175.1 |
| Firmicutes | Bacilli | Lactobacillales | Streptococcaceae | Streptococcus | Streptococcus_oralis | GCF_016549395.1 |
| Firmicutes | Bacilli | Lactobacillales | Streptococcaceae | Streptococcus | Streptococcus_mitis | GCF_016658865.1 |
| Proteobacteria | Betaproteobacteria | Neisseriales | Neisseriaceae | Neisseria | Neisseria_mucosa | GCF_000186165.1 |
| Proteobacteria | Betaproteobacteria | Neisseriales | Neisseriaceae | Neisseria | Neisseria_sicca | GCF_000193735.1 |
| Proteobacteria | Gammaproteobacteria | Pasteurellales | Pasteurellaceae | Haemophilus | Haemophilus_parainfluenzae | GCF_003390455.1 |
| Fusobacteria | Fusobacteriia | Fusobacteriales | Leptotrichiaceae | Leptotrichia | Leptotrichia_wadei | GCF_007990545.2 |
| Fusobacteria | Fusobacteriia | Fusobacteriales | Fusobacteriaceae | Fusobacterium | Fusobacterium_nucleatum | GCF_000158275.2 |
| Firmicutes | Bacilli | Lactobacillales | Carnobacteriaceae | Granulicatella | Granulicatella_elegans | GCF_000162475.2 |
| Firmicutes | Clostridia | Clostridiales | Lachnospiraceae | Lachnoanaerobaculum | Lachnoanaerobaculum_saburreum | GCF_000185385.1 |
| Bacteroidetes | Flavobacteriia | Flavobacteriales | Flavobacteriaceae | Capnocytophaga | Capnocytophaga_leadbetteri | GCF_002302615.1 |
| Firmicutes | Bacilli | Lactobacillales | Streptococcaceae | Streptococcus | Streptococcus_mutans | GCF_008831325.1 |
| Actinobacteria | Actinobacteria | Actinomycetales | Actinomycetaceae | Actinomyces | Actinomyces_oris | GCF_016127955.1 |
| Proteobacteria | Betaproteobacteria | Neisseriales | Neisseriaceae | Neisseria | Neisseria_sp_oral_taxon_014 | GCF_000090875.1 |
| Actinobacteria | Actinobacteria | Micrococcales | Micrococcaceae | Rothia | Rothia_dentocariosa | GCF_000164695.2 |
| Bacteroidetes | Flavobacteriia | Flavobacteriales | Flavobacteriaceae | Capnocytophaga | Capnocytophaga_sputigena | GCF_000173675.1 |
| Bacteroidetes | Flavobacteriia | Flavobacteriales | Flavobacteriaceae | Capnocytophaga | Capnocytophaga_gingivalis | GCF_000174755.1 |
| Proteobacteria | Gammaproteobacteria | Pasteurellales | Pasteurellaceae | Aggregatibacter | Aggregatibacter_sp_oral_taxon_458 | GCF_000466335.1 |
| Firmicutes | Bacilli | Lactobacillales | Streptococcaceae | Streptococcus | Streptococcus_anginosus_group | GCF_001412635.1 |
| Actinobacteria | Actinobacteria | Actinomycetales | Actinomycetaceae | Actinobaculum | Actinobaculum_sp_oral_taxon_183 | GCF_003073475.1 |
| Proteobacteria | Betaproteobacteria | Neisseriales | Neisseriaceae | Neisseria | Neisseria_elongata | GCF_003351685.1 |
| Actinobacteria | Actinobacteria | Micrococcales | Micrococcaceae | Rothia | Rothia_aeria | GCF_016726365.1 |

#### Table S4

**dRNA-seq datasets curated in this study**

| Published year | Species | Phylum | Reference genome | PMID |
|---|---|---|---|---|
| 2011 | *Xanthomonas campestris pv. campestrie* B100 | *Proteobacteria* | GCF_000009165.1 | 22080557 |
| 2012 | *Listeria monocytogenes EGD-e* | *Firmicutes* | GCF_000196035.1 | 22617957 |
| 2012 | *Salmonella enterica subsp. enterica serovar Typhimurium str. SL1344* | *Proteobacteria* | GCF_000210855.2 | 22538806 |
| 2013 | *Corynebacterium glutamicum* ATCC 13032 | *Actinobacteria* | GCF_000011325.1 | 24341750 |
| 2013 | *Campylobacter jejuni 81116* | *Campylobacterota* | GCF_000017905.1 | 23696746 |
| 2014 | *Caulobacter crescentus* | *Proteobacteria* | GCF_000022005.1 | 25078267 |
| 2017 | *Borrelia burgdorferi* | *Spirochaetota* | GCF_000008685.2 | 27913725 |
| 2018 | *Synechococcus elongatus* UTEX 2973 | *Cyanobacteria* | GCF_000817325.1 | 30127850 |
| 2020 | *Bacteroides thetaiotaomicron* VPI-5482 | *Bacteroidota* | GCF_000011065.1 | 32678091 |
| 2020 | *Enterococcus faecalis* | *Firmicutes* | GCF_000007785.1 | 33324581 |
| 2021 | *Flavobacterium psychrophilum* | *Bacteroidota* | GCF_900130145.1 | 36739365 |
| 2021 | *Nostoc punctiforme ATCC* 29133 | *Cyanobacteria* | GCF_000020025.1 | 34779764 |
| 2021 | *Clostridioides difficile* | *Firmicutes* | GCF_000932055.2 | 34131082 |
| 2021 | *Fusobacterium nucleatum subsp. nucleatum* ATCC 25586 | *Fusobacteriota* | GCF_000007325.1 | 34239075 |

**Table S5**

**Curated *Cyanobacteria* traits**

| refseq id | species | unicellular | marine | nitrogen fixation | FaRLiP | LoLiP | RDT fraction |
|---|---|---|---|---|---|---|---|
| GCF_000317675.1 | *Cyanobacterium aponinum PCC 10605* | 1 | 0 | 0 | 0 | 0 | 0.229455 |
| GCF_000316645.1 | *Nostoc sp. PCC 7524* | 0 | 0 | 0 | 0 | 0 | 0.212723 |
| GCF_000317125.1 | *Chroococcidiopsis thermalis PCC 7203* | 0 | 0 | 1 | 1 | 1 | 0.180771 |
| GCF_000317225.1 | *Fischerella thermalis PCC 7521* | 0 | 0 | 1 | 1 | 0 | 0.177499 |
| GCF_001548455.1 | *Fischerella sp. NIES-3754* | 0 | 0 | 1 | 1 | 0 | 0.170573 |
| GCF_000009705.1 | *Nostoc sp. PCC 7120* | 0 | 0 | 1 | 0 | 0 | 0.166333 |
| GCF_000204075.1 | *Anabaena variabilis ATCC 29413* | 0 | 0 | 1 | 0 | 0 | 0.160538 |
| GCF_000316665.1 | *Rivularia sp. PCC 7116* | 0 | 1 | 1 | 0 | 0 | 0.157176 |
| GCF_000010625.1 | *Microcystis aeruginosa NIES-843* | 1 | 0 | 0 | 0 | 0 | 0.14822 |
| GCF_000317475.1 | *Oscillatoria nigro-viridis PCC 7112* | 0 | 0 | 0 | 0 | 0 | 0.146976 |
| GCF_000317515.1 | *Microcoleus sp. PCC 7113* | 0 | 0 | 1 | 0 | 0 | 0.144814 |
| GCF_000317555.1 | *Gloeocapsa sp. PCC 7428* | 0 | 0 | 0 | 0 | 1 | 0.143608 |
| GCF_000020025.1 | *Nostoc punctiforme PCC 73102* | 0 | 0 | 1 | 0 | 0 | 0.142439 |
| GCF_003990575.1 | *Chlorogloeopsis fritschii PCC 6912* | 0 | 0 | 1 | 1 | 1 | 0.13974 |
| GCF_000734895.2 | *Calothrix sp. 336/3* | 0 | 0 | 1 | 0 | 0 | 0.138355 |
| GCF_000196515.1 | *Nostoc azollae 0708* | 0 | 0 | 1 | 0 | 0 | 0.133217 |
| GCF_001277295.1 | *Anabaena sp. wa102* | 0 | 0 | 1 | 0 | 0 | 0.130879 |
| GCF_001264245.1 | *Microcystis panniformis FACHB-1757* | 1 | 0 | 0 | 0 | 0 | 0.127854 |
| GCF_000317695.1 | *Anabaena cylindrica PCC 7122* | 0 | 0 | 1 | 0 | 0 | 0.122553 |
| GCF_000316515.1 | *Cyanobium gracile PCC 6307* | 1 | 0 | 0 | 0 | 0 | 0.119678 |
| GCF_000316575.1 | *Calothrix sp. PCC 7507* | 0 | 0 | 1 | 1 | 0 | 0.115802 |
| GCF_000317025.1 | *Pleurocapsa sp. PCC 7327* | 0 | 0 | 1 | 1 | 0 | 0.110542 |
| GCF_000312705.1 | *Anabaena sp. 90* | 0 | 0 | 1 | 0 | 0 | 0.109736 |
| GCF_000316685.1 | *Synechococcus sp. PCC 6312* | 1 | 0 | 0 | 0 | 0 | 0.106557 |
| GCF_000317105.1 | *Oscillatoria acuminata PCC 6304* | 0 | 0 | 0 | 0 | 0 | 0.105058 |
| GCF_000015705.1 | *Prochlorococcus marinus str. MIT 9303* | 1 | 1 | 0 | 0 | 0 | 0.103865 |
| GCF_000007925.1 | *Prochlorococcus marinus subsp. marinus str. CCMP1375* | 1 | 1 | 0 | 0 | 0 | 0.099237 |
| GCF_000021805.1 | *Cyanothece sp. PCC 8801* | 1 | 0 | 1 | 0 | 0 | 0.097436 |
| GCF_000017845.1 | *Cyanothece sp. ATCC 51142* | 1 | 1 | 1 | 0 | 0 | 0.096753 |
| GCF_000011485.1 | *Prochlorococcus marinus str. MIT 9313* | 1 | 1 | 0 | 0 | 0 | 0.094851 |
| GCF_000317435.1 | *Calothrix sp. PCC 6303* | 0 | 0 | 1 | 0 | 0 | 0.092162 |
| GCF_000316625.1 | *Nostoc sp. PCC 7107* | 0 | 0 | 1 | 0 | 0 | 0.091164 |
| GCF_000317615.1 | *Dactylococcopsis salina PCC 8305* | 1 | 1 | 0 | 0 | 0 | 0.088571 |
| GCF_000155595.1 | *Synechococcus sp. PCC 7335* | 1 | 1 | 1 | 1 | 1 | 0.087398 |
| GCF_000317495.1 | *Crinalium epipsammum PCC 9333* | 0 | 0 | 0 | 0 | 0 | 0.085963 |
| GCF_000014585.1 | *Synechococcus sp. CC9311* | 1 | 1 | 0 | 0 | 0 | 0.081871 |
| GCF_000484535.1 | *Gloeobacter kilaueensis JS1* | 1 | 0 | 0 | 0 | 0 | 0.081865 |
| GCF_000270265.1 | *Synechocystis sp. PCC 6803 substr. GT-S* | 1 | 0 | 0 | 0 | 0 | 0.080893 |
| GCF_000012505.1 | *Synechococcus sp. CC9902* | 1 | 1 | 0 | 0 | 0 | 0.080357 |
| GCF_000317635.1 | *Halothece sp. PCC 7418* | 1 | 1 | 1 | 0 | 0 | 0.076628 |
| GCF_000018105.1 | *Acaryochloris marina MBIC11017* | 1 | 1 | 0 | 0 | 0 | 0.076616 |
| GCF_000317065.1 | *Pseudanabaena sp. PCC 7367* | 0 | 1 | 0 | 0 | 0 | 0.074232 |
| GCF_000021825.1 | *Cyanothece sp. PCC 7424* | 1 | 0 | 1 | 0 | 0 | 0.071579 |

| | | | | | | | |
|---|---|---|---|---|---|---|---|
| GCF_000317045.1 | *Geitlerinema sp. PCC 7407* | 0 | 0 | 0 | 0 | 0 | 0.070746 |
| GCF_000011385.1 | *Gloeobacter violaceus PCC 7421* | 1 | 0 | 0 | 0 | 0 | 0.069853 |
| GCF_000284135.1 | *Synechocystis sp. PCC 6803 substr. GT-I* | 1 | 0 | 0 | 0 | 0 | 0.069307 |
| GCF_000284215.1 | *Synechocystis sp. PCC 6803 substr. PCC-N* | 1 | 0 | 0 | 0 | 0 | 0.068966 |
| GCF_000147335.1 | *Cyanothece sp. PCC 7822* | 1 | 0 | 1 | 0 | 0 | 0.068219 |
| GCF_000195975.1 | *Synechococcus sp. WH 8102* | 1 | 1 | 0 | 0 | 0 | 0.066158 |
| GCF_000009725.1 | *Synechocystis sp. PCC 6803* | 1 | 0 | 0 | 0 | 0 | 0.064767 |
| GCF_000013205.1 | *Synechococcus sp. JA-3-3Ab* | 1 | 0 | 1 | 0 | 0 | 0.060478 |
| GCF_000317575.1 | *Stanieria cyanosphaera PCC 7437* | 0 | 0 | 0 | 0 | 0 | 0.060218 |
| GCF_000013225.1 | *Synechococcus sp. JA-2-3B'a(2-13)* | 1 | 0 | 1 | 0 | 0 | 0.05814 |
| GCF_000316605.1 | *Leptolyngbya sp. PCC 7376* | 0 | 1 | 0 | 0 | 0 | 0.057924 |
| GCF_000161795.2 | *Synechococcus sp. WH 8109* | 1 | 1 | 0 | 0 | 0 | 0.056206 |
| GCF_000015645.1 | *Prochlorococcus marinus str. AS9601* | 1 | 1 | 0 | 0 | 0 | 0.054054 |
| GCF_000012625.1 | *Synechococcus sp. CC9605* | 1 | 1 | 0 | 0 | 0 | 0.050781 |
| GCF_000012465.1 | *Prochlorococcus marinus str. NATL2A* | 1 | 1 | 0 | 0 | 0 | 0.049563 |
| GCF_000015685.1 | *Prochlorococcus marinus str. NATL1A* | 1 | 1 | 0 | 0 | 0 | 0.047478 |
| GCF_000011465.1 | *Prochlorococcus marinus subsp. pastoris str. CCMP1986* | 1 | 1 | 0 | 0 | 0 | 0.047431 |
| GCF_000012525.1 | *Synechococcus elongatus PCC 7942* | 1 | 0 | 0 | 0 | 0 | 0.046729 |
| GCF_000019485.1 | *Synechococcus sp. PCC 7002* | 1 | 0 | 0 | 0 | 0 | 0.043742 |
| GCF_000015665.1 | *Prochlorococcus marinus str. MIT 9515* | 1 | 1 | 0 | 0 | 0 | 0.042735 |
| GCF_000010065.1 | *Synechococcus elongatus PCC 6301* | 1 | 0 | 0 | 0 | 0 | 0.040193 |
| GCF_000011345.1 | *Thermosynechococcus elongatus BP-1* | 1 | 0 | 0 | 0 | 0 | 0.035714 |
| GCF_000018065.1 | *Prochlorococcus marinus str. MIT 9215* | 1 | 1 | 0 | 0 | 0 | 0.033582 |
| GCF_000015965.1 | *Prochlorococcus marinus str. MIT 9301* | 1 | 1 | 0 | 0 | 0 | 0.027778 |
| GCF_000012645.1 | *Prochlorococcus marinus str. MIT 9312* | 1 | 1 | 0 | 0 | 0 | 0.021834 |

#### Table S6

**Recombinant plasmids constructed for RUT test with the fluorescence reporter assay**

| Plasmids | Putative RUTs | Putative RUT sequences(5'→3') |
|---|---|---|
| pXG10SFM | / | / |
| pXG10SFM0 | / | / |
| pXG10SFM-7120RUT1 | nifD\|nifH | ATTCCTCTTCCCACTCTCCCTTCCCGACTCCTCACTCTCCCAAATATACTTCTATTCCCCCA |
| pXG10SFM-7120RUT2 | nifD-2 | TGTTCTTTTCCCTCTCAACCGTGGTGTTTCCATTCAATCT |
| pXG10SFM-7120RUT3 | nifK-1 | TCCTCTCTTCGATCGCCACCACTTACACCGCTATTCTACCCTCGGCTACCAAGGTGGTCT |
| pXG10SFM-7120RUT4 | nifK-2 | ACTTCCGTTCCTTGTTGTTCACCGAGCCTGTAGACTTCTTCATCGGTAACTCCT |
| pXG10SFM-7120RUT5 | nifK-3 | GTTCGCTATCTACGGCGATCCAGATTTGATCATCTCCATCACCAGCTTCT |
| pXG10SFM-7120RUT6 | nifK-4 | CGCGTTCATCACCAACTCCAAGAACGCTGGTTCTATTCCTCAAGATTTCCCCGTACCCT |
| pXG10SFM-7120RUT7 | nifK-5 | TGCTTACTTCCGTACACACCTCAGCCGTCACTACAAAGAGCCTTGCTCCGCAGTATCTTCT |
| pXG10SFM-6912RUT1 | rfpB\|psbA4 | TTACAGAACCCCGACTTCTCCAAGAAGTCAAGGATCTC |
| pXG10SFM-6912RUT2 | rfpA-1 | CGCTCAAGCCATCCTCTTAGAGCAAAGCCGCCTTCAAGCCCAACAAGAAGCCACAATTAATCG |
| pXG10SFM-6912RUT3 | rfpA-2 | AACAATCTTCCCCCAATTCCCCAGTTCCCGATCTCCTCGAAGAACATCC |
| pXG10SFM-6912RUT4 | rfpA-3 | CCACTTTGCCATGGCAATTCATCAATATCAACTGTATCAGCAAGTTCA |
| pXG10SFM-6912RUT5 | psbA3-1 | ACAGCACCTTCCATCACTGCCTAGTTGGTTTTCAGTTGTCAGTTTGATTCCTCCCTGACAACTGAACCATTTCC |
| pXG10SFM-6912RUT6 | psbA3-2 | GATAAATTGTCGGTTTTAACCCAAATATCATCAACAAAACCCGCCCCTACTAACTAAAAAATTC |
| pXG10SFM-6912RUT7 | isiX | TTACGGCCCCACATTACATCTCAAGTCATCTCTATTACCTTACTACTTCGATCCCGCCCA |
| pXG10SFM-6912RUT8 | apcB3 | CGCAAAATACCTCTCTAACTCTTATTCCTCCGTGTCCTTTGTATCTCTGTGAAAAAAACGAACC |

**Table S7**

#### Primers used in this study

| Primer | Sequence(5'→3') |
|---|---|
| SFM0-F | TGGATGAGCTCTACAAATAATCTAGAACCAGTAACGTTATACGATG |
| SFM0-R | GTTAAACAAAATTATTTCTAGGgtacCGTTTTCGCAGAAACGTG |
| 7120RUT1-F | TGGATGAGCTCTACAAATAATCTAGAATTCCTCTTCCCACTCTC |
| 7120RUT1-R | ACAAAATTATTTCTAGGgtacCTGGGGGAATAGAAGTATATTTG |
| 7120RUT2-F | TGGATGAGCTCTACAAATAATCTAGATGTTCTTTTCCCTCTCAAC |
| 7120RUT2-R | ACAAAATTATTTCTAGGgtacCAGATTGAATGGAAACACCAC |
| 7120RUT3-F | TGGATGAGCTCTACAAATAATCTAGATCCTCTCTTCGATCGCCAC |
| 7120RUT3-R | ACAAAATTATTTCTAGGgtacCAGACCACCTTGGTAGCCGAG |
| 7120RUT4-F | TGGATGAGCTCTACAAATAATCTAGAACTTCCGTTCCTTGTTGTTC |
| 7120RUT4-R | ACAAAATTATTTCTAGGgtacCAGGAGTTACCGATGAAGAAG |
| 7120RUT5-F | TGGATGAGCTCTACAAATAATCTAGAGTTCGCTATCTACGGCGATC |
| 7120RUT5-R | ACAAAATTATTTCTAGGgtacCAGAAGCTGGTGATGGAG |
| 7120RUT6-F | TGGATGAGCTCTACAAATAATCTAGACGCGTTCATCACCAACTC |
| 7120RUT6-R | ACAAAATTATTTCTAGGgtacCAGGGTACGGGGAAATCTTG |
| 7120RUT7-F | TGGATGAGCTCTACAAATAATCTAGATGCTTACTTCCGTACACAC |
| 7120RUT7-R | ACAAAATTATTTCTAGGgtacCAGAAGATACTGCGGAGCAAG |
| 6912RUT1-F | TGGATGAGCTCTACAAATAATCTAGATTACAGAACCCCGACTTCTC |
| 6912RUT1-R | GTTAAACAAAATTATTTCTAGGgtacCGAGATCCTTGACTTCTTGGAG |
| 6912RUT2-F | TGGATGAGCTCTACAAATAATCTAGACGCTCAAGCCATCCTCTTAG |
| 6912RUT2-R | GTTAAACAAAATTATTTCTAGGgtacCCGATTAATTGTGGCTTCTTG |
| 6912RUT3-F | TGGATGAGCTCTACAAATAATCTAGAAACAATCTTCCCCCAATTCC |
| 6912RUT3-R | GTTAAACAAAATTATTTCTAGGgtacCGGATGTTCTTCGAGGAGATC |
| 6912RUT4-F | TGGATGAGCTCTACAAATAATCTAGACCACTTTGCCATGGCAATTC |
| 6912RUT4-R | GTTAAACAAAATTATTTCTAGGgtacCTGAACTTGCTGATACAGTTG |
| 6912RUT5-F | TGGATGAGCTCTACAAATAATCTAGAACAGCACCTTCCATCAC |
| 6912RUT5-R | GTTAAACAAAATTATTTCTAGGgtacCGGAAATGGTTCAGTTGTCAG |
| 6912RUT6-F | TGGATGAGCTCTACAAATAATCTAGAGATAAATTGTCGGTTTTAAC |
| 6912RUT6-R | CTTAAAGTTAAACAAAATTATTTCTAGGgtacCGAATTTTTTAGTTAGTAGG |
| 6912RUT7-F | TGGATGAGCTCTACAAATAATCTAGATTACGGCCCCACATTACATC |
| 6912RUT7-R | GTTAAACAAAATTATTTCTAGGgtacCTGGGCGGGATCGAAGTAG |
| 6912RUT8-F | TGGATGAGCTCTACAAATAATCTAGACGCAAAATACCTCTCTAACTC |
| 6912RUT8-R | GTTAAACAAAATTATTTCTAGGgtacCGGTTCGTTTTTTTCACAGAG |

### Supplementary Dataset

**Dataset S1**. Protein coding gene clusters and Rfam families that were used for data augmentation.
